# Cryptic bacterial pathogens of diatoms peak during senescence of a winter diatom bloom

**DOI:** 10.1101/2023.06.22.545670

**Authors:** Laura Branscombe, Ellen L. Harrison, Choong Zhi Yi Daniel, Matthew Keys, Claire Widdicombe, William H. Wilson, Michael Cunliffe, Katherine Helliwell

## Abstract

Diatoms are globally abundant algae that form extensive blooms in aquatic ecosystems. Certain bacteria behave antagonistically towards diatoms, killing or inhibiting their growth. Despite their crucial implications to diatom health and bloom control, insight of the prevalence and dynamics of antagonistic bacteria in nature is lacking. We report an ecosystem assessment of the diversity and seasonal patterns of bacterial antagonists of diatoms via regular plaque-assay sampling in the Western English Channel (WEC), where diatoms frequently bloom. Unexpectedly, peaks in antagonist detection did not occur during characteristic spring blooms, but coincided with a winter bloom of *Coscinodiscus*, suggesting bacterial pathogens likely influence distinct diatom host populations. We isolated multiple antagonists, spanning 4 classes and 10 bacterial orders. Many species had no prior reports of pathogenicity towards diatoms, and we verified diatom growth inhibitory effects of 8 isolates. In all cases tested, pathogenicity was activated by pre-exposure to diatom organic matter. Discovery of widespread ‘cryptic’ antagonistic activity evident under specific conditions, indicates that bacterial pathogenicity towards diatoms is more prevalent than previously recognised. Finally, mining *Tara* Oceans data revealed the global biogeography of WEC antagonists and co-occurrence patterns with diatom hosts. Our study indicates that multiple, diverse antagonistic bacteria have potential to impact diatom growth and bloom dynamics in marine waters globally.

## Introduction

Diatoms are single-celled photosynthetic eukaryotes that contribute around 40% of marine primary productivity (1). Diatoms frequently form large, sometimes toxic blooms, during which they can constitute up to 90% of the phytoplankton community (2). Organic matter produced during a bloom is vital for sustaining planktonic life (3). Biomass that is not recycled can form nutrient-rich aggregates that sink to the sea floor, sequestering carbon in the deep ocean (4). It is thus crucial to understand environmental drivers controlling diatom bloom dynamics. A range of abiotic (e.g. nutrients, light, and temperature) and biotic (e.g. grazers and viruses) factors control bloom formation and demise (3). However, the relative contribution of these drivers, and the spectrum of biotic pressures influencing phytoplankton populations in the ocean, remain open to debate (3). Given that around half of the carbon that is fixed by phytoplankton such as diatoms is processed by bacteria (5,6), gaining insight of the interactions between diatoms and bacteria and their influence on natural bloom dynamics is crucial for better understanding nutrient fluxes and the fate of carbon in the oceans.

Diatoms engage in a suite of biotic interactions with bacteria, ranging from synergistic to antagonistic (7,8). Accumulating evidence indicates that diatoms host species-specific bacterial communities, a so-called ‘diatom microbiome’ (8,9). Bacterial taxa typically belonging to *Alphaproteobacteria*, *Bacteroidetes* and *Gammaproteobacteria*, are frequently associated with diatoms (10,11), with certain genera in particular (e.g., *Sulfitobacter*, *Roseobacter*, *Alteromonas*, and *Flavobacterium*) most commonly identified (8). Notably, several interactions between diatoms and members of the *Rhodobacteraceae* (of the *Alphaproteobacteria*) are synergistic (12–15), with bacteria conferring protection against environmental stressors (e.g. oxidative stress (15)) and/or alleviating the demands of diatoms for nutrients such as vitamins 1. (12) and nitrogen (13,14).

Certain bacteria can also have algicidal (causing cell death) and/or algistatic (growth inhibitory) effects against diatoms (7,16). These antagonistic bacteria tend to belong to the *Gammaproteobacteria* (17), specifically of the orders *Alteromonadales* and *Pseudomonadales*, as well as *Flavobacteria* (7). Current understanding of the mechanistic basis of antagonistic diatom-bacteria interactions has been yielded largely through study of culture-based model systems. For instance, the Flavobacterium *Kordia algicida* displays algicidal activities against the diatoms *Thalassiosira weissflogii*, *Phaeodactylum tricornutum* and *Skeletonema costatum* (18), via release of extracellular proteases that cause diatom cell lysis (18). Another Flavobacterium, *Croceibacter atlanticus,* is also capable of inhibiting growth of various diatoms, but to different degrees (e.g. *Pseudo-nitzschia multiseries* PC9 by 73%, and *Thalassiosira pseudonana* by 33%) (19). Laboratory study demonstrated that *C. atlanticus* directly attaches to diatoms and inhibits cell division. Treatment of *T. pseudonana* with *C. atlanticus* exudates resulted in increases in intra- and extra-cellular carbon, suggesting that bacterial antagonists can modulate host metabolism to support their growth (20).

Whilst model systems have been instrumental in elucidating impacts of algicidal bacteria on diatom physiology, insights of the prevalence and dynamics of diatom algicidal bacteria in nature is fundamentally lacking. This is in part due to sampling constraints, with strains typically being identified from non-axenic algal cultures (13,19) or collected ad-hoc from different geographical locations (17). Moreover, in general there has been a bias in sampling efforts towards the study of algicidal bacteria of dinoflagellates, which are of particular interest due to their potential for controlling harmful dinoflagellate blooms (17). These factors have precluded ecosystem-specific trends being drawn on the population structure and seasonal patterns of diatom antagonistic bacteria, and many open questions remain. For instance, the prevalence and diversity of bacterial antagonists that co-occur with diatoms in the same ecosystem remains unclear. This, coupled with knowledge of the host range of antagonists towards diatoms and host susceptibility to different bacteria inhabiting the same ecosystem, is necessary to better understand the ecological relevance of bacterial antagonists. Gaining insights of their seasonal dynamics is also necessary to explore co-occurrences with host species. Finally, we have little knowledge of how broadly distributed bacterial antagonists are in marine ecosystems globally. To address these questions a more systematic ecosystem approach is necessary to characterise the bacterial pathobiome of diatoms in nature.

The Western Channel Observatory (WCO) represents one of the most comprehensive oceanographic time series in the world, collecting plankton abundance data weekly throughout the year (21). Diatoms can dominate Western English Channel (WEC) phytoplankton communities, with spring blooms frequently made up of *Chaetoceros*, *Thalassiosira* and *Skeletonema* (21). Peaks of certain diatoms during winter months are also well documented (22,23). WEC bacterial communities also exhibit robust seasonal patterns (24,25). *Alphaproteobacteria*, namely *Rickettsiales* (SAR11) and *Rhodobacterales*, constitute the most abundant taxa, in addition to *Flavobacteriales* (*Bacteroidetes*), and *Gammaproteobacteria* (*Vibrionales* and *Pseudomonadales*) (25). Notably, abundances of *Flavobacterale* and *Rhodobacterale* populations correlated significantly with phytoplankton-derived polysaccharides during a spring WEC diatom bloom (26), suggesting tight coupling of bacterial population structure and bloom succession. The accessibility of the WEC off the coast of Plymouth lends itself to routine studies that have been invaluable for the identification of interactions between planktonic groups. For instance, relationships between WEC diatoms and mycoplankton have been revealed through detection of concurrent diatom and chytrid blooms observed interannually in the WEC (27). Additionally, single-cell picking approaches led to the isolation of a novel thraustochytrid ‘ThrauL4’ that parasitizes *Chaetoceros* (28).

These studies highlight the power of long-term monitoring coupled with environmental sampling approaches that ‘bring-into-the-lab’ biotic associations representative of those in nature. Here, we exploit our unique access to WEC microbial communities to systematically assess the diversity of bacterial antagonists of diatoms in the WEC. Utilising a plaque assay approach (29) over the course of an annual cycle, we have characterised the diversity and physiological traits of the bacterial ‘pathobiome’ of bloom-forming diatoms residing in the WEC. Our study uncovers seasonal patterns of previously unreported antagonistic activity against diatoms in a diverse range of bacterial lineages.

## Results

### A systematic environmental pipeline to isolate naturally occurring diatom antagonists

Exploiting routine access to WEC microbial communities, we developed a systematic environmental sampling pipeline to isolate diatom pathogens, employing a soft agar overlay plaque assay technique. This entailed inoculating concentrated suspensions of five diatom hosts with the bacterial fraction of WEC seawater (Methods; Figure S1A), with appearance of regions of clearance or ‘plaques’ on the diatom lawns indicative of diatom growth inhibition. Diatoms chosen for this study included four centric diatoms: *T. pseudonana* (PLY693), *T. weissflogii* (PLY541), *Skeletonema* sp. PLY627, and *Chaetoceros* sp. PLY617, alongside the model pennate *P. tricornutum* (PLY100) (Table S1). Several of these cultures were originally isolated from the WEC, and all are documented to inhabit WEC waters, with *Thalassiosira*, *Skeletonema* and *Chaetoceros* representing some of the most abundant diatom genera in the WEC (21) and indeed globally (30). Whilst we never observed plaques on control assay plates (plaque assays inoculated with autoclaved seawater), plaques were frequently observed on assays inoculated with the bacterial fraction of seawater (Figure 1A). In total, across the five diatom hosts and over 13 months of sampling, we observed 181 plaques. The greatest number of plaques were obtained on planktonic centric diatoms, with *T. pseudonana* being the most ‘susceptible’ host having 56 plaques observed in total, followed by *T. weissflogii* (53 plaques), *Skeletonema* sp. PL7627 (44 plaques), and *Chaetoceros* sp. PLY617 (23 plaques) (Figure 1B). By comparison, only 5 plaques were isolated on the benthic pennate *P. tricornutum*. The number of plaques observed per month also varied, with values ranging from none (Nov 2020 and May 2021) to over a hundred across the 5 diatom hosts (Dec 2020) (Figure 1C).

**Figure 1.**
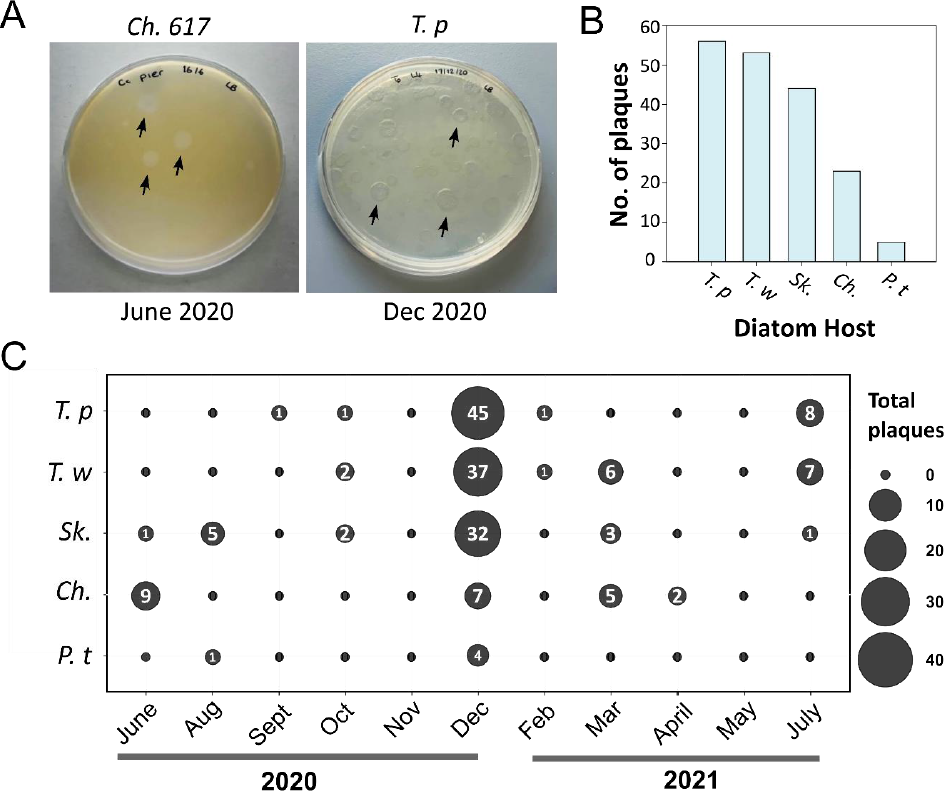
Systematic isolation of naturally occurring diatom antagonists in the Western English Channel. **A.** Photographs of an example *Chaetoceros* sp. PLY617 and *T. pseudonana* plaque assay plate from June 2020 and Dec 2020, respectively. Several clearance plaques are indicated (black arrows). **B.** Total number of plaques obtained for each diatom host over the 13-month sampling period. **C.** Bubble plot showing the number of plaques observed per diatom host each sampling month.

### Recurrent isolation of diverse WEC plaque-forming bacteria spanning 4 classes and 10 orders of bacterial diversity

Bacteria isolated from individual plaques were identified via 16S rRNA gene sequencing (Methods). This was done for each plaque obtained during the sampling, with the exception of Dec 2020, for which three plaques per host were randomly selected, as the number of plaques at this time point (125) was too great to process. In a total of 65 plaques identified, 18 different bacterial species were identified, spanning 4 classes and 10 orders of bacteria (Figure S2; Dataset S1). Bacteria belonging to the *Alphaproteobacteria* were most frequently identified (over 50% of plaques), followed by *Gammaproteobacteria* (40.8%), *Flavobacteria* (4.2%), and least frequently were bacterial species belonging to the class *Bacilli* (1.4%) (Figure 2A). Examining bacterial class by host, we found that *Alphaproteobacteria* were most frequently isolated from plaque assays with the centric diatoms *Skeletonema* sp. PLY627 and *Chaetoceros* sp. PLY617 (Figure 2B). By comparison, *Gammaproteobacteria* were detected least frequently on *Skeletonema* sp. PLY627, being detected predominantly on the *Thalassiosira* species alongside *Chaetoceros* sp. PLY617. Notably, certain bacterial species were isolated on multiple independent occasions. For instance, the *Roseobacter Ponticoccus alexandrii* (*Alphaproteobacteria*) was detected on 19 independent occasions on 7 sampling months, and across all four centric diatom hosts, but not *P. tricornutum* (Figure 2C-D). The *Gammaproteobacteria H. titanicae* and *M. adhaerens* were detected the second and third most frequently, being identified in 13 and 8 plaques, respectively.

**Figure 2.**
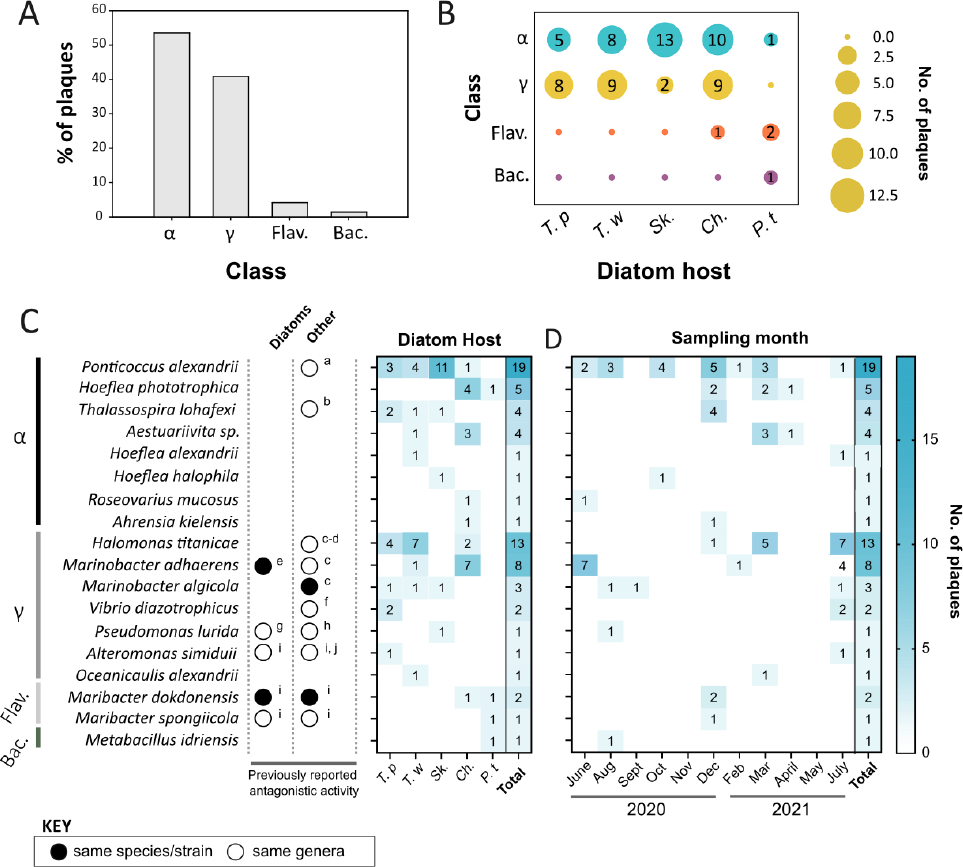
Recurrent isolation of diverse bacteria from plaque assays on different diatom hosts and sampling months. **A.** Percentage of plaques identified to contain bacteria belonging to the classes *Alphaproteobacteria* (α), *Gammaproteobacteria* (γ), *Flavobacteria* (Flav.) and *Bacilli* (Bac.), respectively. **B.** Bubble plot showing the number of plaques identified containing bacteria belonging to each bacterial class per diatom host species. C. Heat map showing the no. of plaques for each WEC bacterial isolate across different diatom hosts. Those species that have previously been reported to confer antagonistic activity against diatoms and/or other eukaryotic phytoplankton are indicated, with the relevant literature cited as follows: ^a^(31), ^b^(32), ^c^(33), ^d^(64), ^e^(34), ^f^(35), ^g^(37), ^h^(36), ^i^(39), ^j^(38). The bacterial classes are also labelled. D. Heatmap of no. of plaques identified per bacterium each sampling month. N.b. For the Dec 2020 sampling point there were more plaques (125) than could be identified.

To determine whether any of our isolates had prior reports of being algicidal, we conducted a literature review. This revealed that algicidal activity against algae has been reported previously in 8 of the genera of the WEC isolates, including: *Ponticoccus* (31), *Thalassospira* (32), *Halomonas* (33), *Marinobacter* (33,34), *Vibrio* (35), *Pseudomonas* (36,37), *Alteromonas* (38,39) and *Maribacter* (39) (Figure 2C). This was to species level for three bacteria (*Marinobacter algicola, Marinobacter adhaerens* and *Maribacter dokdonensis*). Of the WEC isolates, prior reports of algicidal behaviour towards diatoms has been reported only for *Pseudomonas* (37), *Alteromonas* (39) species, as well as *M. dokdonensis* (39) and *M. adhaerens* (34).

### Seasonally persistent *P. alexandrii* exhibits facultative algicidal activity against diatoms in a specie-specific manner

Closer inspection of plaques clearly showed clearance of diatom cells (Fig. S3), indicating that the bacterial isolates from plaques likely confer a growth inhibitory and/or algicidal effect towards diatoms. To further investigate the effect of WEC plaque-forming bacteria, we conducted a range of experiments both in plaque assay and liquid culture. To focus these investigations, we examined the most frequently isolated bacterium, *P. alexandrii*. To verify that plaques from which *P. alexandrii* was isolated could recapitulate plaque-formation on diatom plates, we inoculated material from dissected plaques into fresh plaque assays. These ‘plaque-to-plaque’ assays resulted in the formation of numerous new plaques on *T. pseudonana*, *Skeletonema* sp. PLY627, and *Chaetoceros* sp. PLY617, but not *T. weissflogii* or *P. tricornutum* plates (Table S2). Incidentally, we could also propagate plaques containing *M. adhaerens,* and *M. algicola* in this manner. We therefore tested whether pure cultures of *P. alexandrii* could cause plaque-formation. Repeated trials inoculating *P. alexandrii* pre-grown on ½YTSS (Yeast Tryptone Sea Salts) agar plates into *T. pseudonana* plaque assays failed to recapitulate plaque formation. Since the plaque-to-plaque assays successfully yielded plaque formation, we postulated that *P. alexandrii* cells growing on diatom soft-overlay plates are in a different physiological state to cells grown on ½YTSS medium. To test the plaque-forming ability of *P. alexandrii* cells grown in conditions more akin to those within a plaque, we tested *P. alexandrii* cells pre-grown on medium made up of autoclaved diatom cultures, hereafter ‘Dead Diatom Medium’ (DDM). Inoculation of DDM-grown *P. alexandrii* at different densities led to the formation of plaques on *T. pseudonana* soft-overlay plates. In contrast, ½YTSS grown *P. alexandrii* yielded no (or few) plaques, even at equivalent bacterial densities (Figure 3A). These experiments suggest that the capacity of *P. alexandrii* to cause plaque formation on *T. pseudonana* plates is induced by pre-exposure to diatom organic matter.

**Figure 3.**
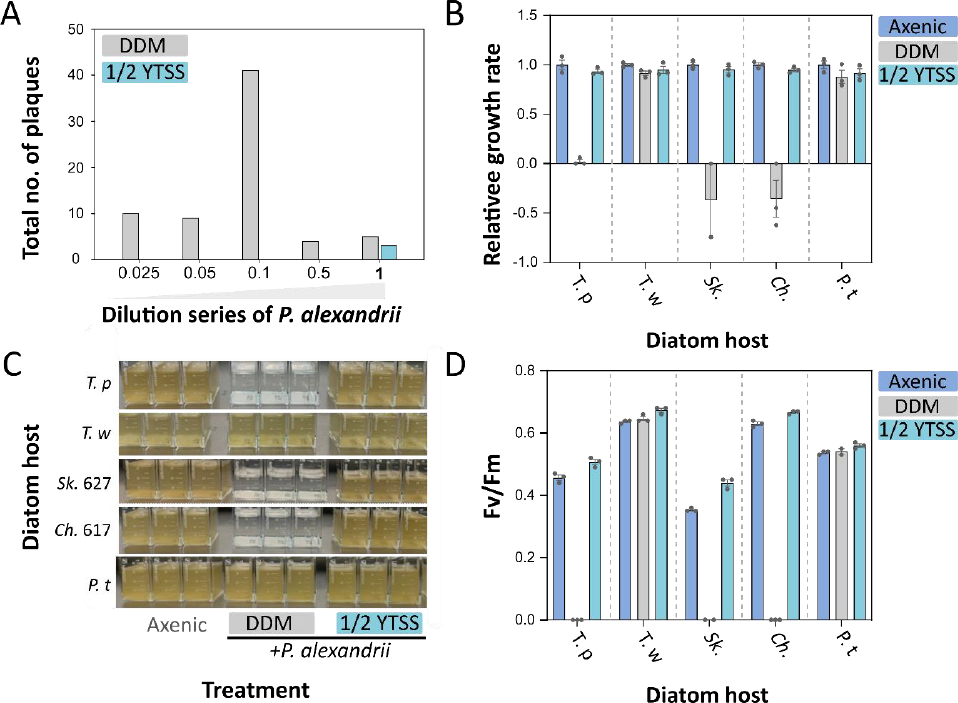
*Ponticoccus alexandrii* inhibits diatom growth in a species-specific manner but only when pre-grown on diatom organic matter. **A.** Number of plaques when pure cultures of *P. alexandrii* were inoculated into plaque assays with *T. pseudonana*, comparing different densities of bacteria. Prior to the plaque assay, *P. alexandrii* was grown either on agar plates made up with autoclaved diatoms (‘Dead Diatom Medium’ (DDM)) or rich ½YTSS (Yeast Tryptone Sea Salts) medium. The 1× treatment represented a final OD600 of 0.01. B. Growth rate relative to axenic control (measured during exponential growth phase) of all five diatom hosts *T. pseudonana* (*T. p*), *T. weissflogii* (*T. w*), *Skeletonema* sp. PLY627 (*Sk.*), *Chaetoceros* sp. PLY617 (*Ch*.), and *P. tricornutum* (Pt) when inoculated with DDM versus ½YTSS grown *P. alexandrii*. Error bars indicate ± S. E. M for n=3; individual data points are plotted. C. Photographs of the cultures described in (B) on day 9. D. Maximum quantum efficiency of photosystem II (Fv/Fm)of cultures on day 8. Error bars indicate ± S. E. M for n=3; individual data points are plotted.

To better characterise this phenomenon, we tested the impact of DDM versus ½YTSS grown *P. alexandrii* on diatom growth in liquid culture. We inoculated diatom hosts into three different treatments: either growing them axenically or co-inoculating them with *P. alexandrii* pre-grown on ½YTSS or DDM. Whereas growth of all five diatoms inoculated with *P. alexandrii* pre-grown on ½YTSS was comparable to the axenic control, growth of *T. pseudonana*, *Skeletonema* sp. PLY627 and *Chaetoceros* sp. (PLY617) was completely impaired by DDM-grown *P. alexandrii* (Figure 3B-C and Figure S4). Similarly, maximum quantum efficiency of photosystem II (Fv/Fm) of *T. pseudonana*, *Skeletonema* sp. PLY627 and *Chaetoceros* sp. PLY617 cells was undetectable when inoculated in co-culture with DDM-grown *P. alexandrii* cells (day 8) (Figure 3D). Therefore, in concurrence with our plaque assay results, these data suggest that the growth-inhibitory effect of *P. alexandrii* on diatoms is facultative, being activated only when cells are first grown on DDM. Finally, we verified that *P. alexandrii* causes diatom cell death, using the live-dead stain SYTOX green. Inoculation of *T. pseudonana* with *P. alexandrii* not only led to reduced growth compared to axenic culture (Figure S5A), but also 95% of cells detected were dead, indicating that this bacterium is algicidal (Figure S5B).

### Activation of antagonistic activity by DDM is widespread amongst diverse WEC antagonists

We extended our approach to determine whether other WEC bacterial strains also exhibit growth inhibitory effects against diatoms, which are activated by pre-exposure to DDM. We tested the impact of inoculating different bacterial isolates into co-culture with the diatom host from which they were originally isolated. We focussed these efforts predominantly on species for which there have been no prior reports of algicidal activity against diatoms. In total, we tested 11 bacterial species (including *P. alexandrii*), including *Thalassospira lohafexi*, *Vibrio diazotrophicus*, *M. adhaerens*, *Halomonas titanicae*, *Metabacillus idriensis*, *Maribacter spongiicola, Hoeflea phototrophica*, *M. algicola*, *Pseudomonas lurida*, as well as *Oceanicaulis alexandrii.* Inoculation of *T. pseudonana* with the *Gammaproteobacteria T. lohafexi* and *V. diazotrophicus* pre-grown on DDM, caused an almost complete (99.8% and 99.7%, respectively) decrease in diatom cell numbers relative to the axenic control (Figure 4A) (*P. alexandrii* was used as a positive control). By comparison, ½YTSS grown *T. lohafexi* and *V. diazotrophicus* cells resulted in a decrease in *T. pseudonana* cell density by 32.3% and 21.7%, respectively. Both *M. adhaerens*, and *H. titanicae* also reduced growth of *Chaetoceros* sp. (PLY617) (causing a 99.8% and 31.8% reduction in diatom density), but only when pre-grown on DDM (Figure 4B). Whilst diatom growth in the DDM-grown *H. titanicae* co-culture was not statistically significantly different compared to the axenic control, it was significantly different (reduced by 35%) compared to the ½YTSS treatment (p-value: <0.05, one-way ANOVA). We also verified the antagonistic effect of *M. idriensis, H. phototrophica and M. spongiicola* that caused a 98.7%, 42.8% and 67.7% reduction in *P. tricornutum* cell numbers when pre-grown on DDM, respectively, compared to the diatom only control (Figure 4C).

**Figure 4.**
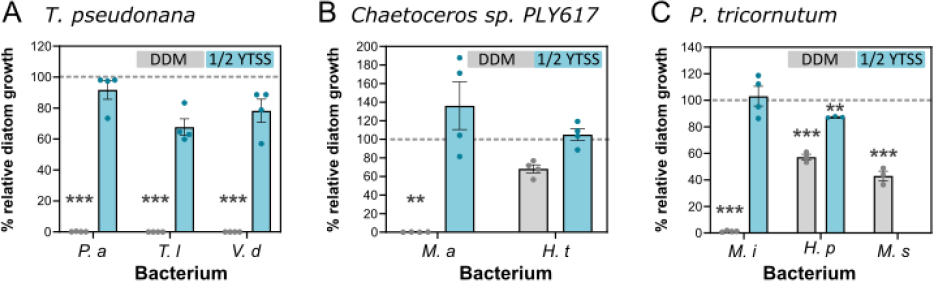
Growth inhibitory effect against diatoms is activated by DDM in multiple other WEC plaque-forming bacteria. Percentage growth relative to axenic control (marked by a dashed grey line) of the diatoms *T. pseudonana* **(A)**, *Chaetoceros* sp. PLY617 **(B)** and *P. tricornutum* **(C)** in the presence of WEC bacteria. Bacterial isolates tested included: *Ponticoccus alexandrii* (*P. a*), *Thalassospira lohafexi* (*T. l*), *Vibrio diazotrophicus* (*V. d*) for *T. pseudonana*; *Marinobacter adhaerens* (*M. a*), and *Halomonas titanicae* (*H. t*) for *Chaetoceros* sp. PLY617, as well as *Metabacillus idriensis (M. i), Hoeflea phototrophica (H. p) and Maribacter spongiicola* (*M. s*) for *P. tricornutum*. N. b. only *M. spongiicola* pre-grown on DDM was tested. The effect on the growth of diatom host on which the bacterium was originally isolated was examined. Error bars indicate ± S. E. M for n=4 or 3; individual data points are plotted. Growth was quantified by measuring diatom cell density (cells/ml) on 4-14 days (depending on the diatom host species) following inoculation into fresh f/2 media with or without WEC bacteria. P-values (one-way ANOVA comparing individual co-culture treatments to the axenic control): P< 0.05; **p < 0.01; ***p < 0.001 (*n*=3 to 4). Individual data points are shown.

However, not all the bacteria isolated via our plaque assay approach could be verified to affect diatom growth in liquid culture. Inoculating *M. algicola* into co-culture with *Skeletonema* sp. PLY627 or *T. weissflogii* did not cause significant reductions in diatom growth in any treatment examined (Figure S6), even though ‘plaque-to-plaque’ assays inoculating from *M. algicola* plaques did lead to formation of multiple new plaques on *T. pseudonana* (Table S2). For *P. lurida* we observed an 11.7% and 21.3% inhibition of diatom host growth (when pre-grown on DDM or ½YTSS, respectively) relative to the axenic control (Figure S6). Similarly, ½YTSS-grown *O. alexandrii* caused a 20.8% reduction of *T. weissflogii* growth, but this was not significant. Thus, under the conditions examined, we have not been able to conclusively verify that these bacteria are true diatom pathogens. However, we have verified that eight (nine including *M. algicola*) of the 11 bacterial isolates tested do confer robust growth inhibitory effects against diatoms (Figure 4). The percentage inhibition of diatom growth by these bacteria ranged from 31.8% (*H. titanicae*) to over 98% (for *P. alexandrii T. lohafexi* and *V. diazotrophicus, M. adhaerens*, and *M. idriensis*), with the effect being greatest when cells were pre-grown on DDM.

### Peaks in antagonistic activity in the WEC coincided with senescence of a winter diatom bloom

Having verified that the plaque assay sampling approach has successfully isolated antagonistic bacteria capable of inhibiting diatom growth, we investigated broader ecosystem trends relating to patterns in plaque enumeration over our sampling. Of particular note, was the large peak in plaques recorded in Dec 2020 (Figure 1C). Examination of diatom diversity and abundance data collected via the WCO, revealed that a winter diatom bloom was in progress in Dec 2020 (Figure 5A). The centric diatom *Coscinodiscus wailesii* accounted for approximately 92% of diatom biomass at the peak of this bloom (on Dec 7^th^), with diatoms making up 83.9% of the total phytoplankton biomass. *Thalassiosira* but not *Skeletonema* and *Chaetoceros* were also present (Figure S7A-B). Notably, images of phytoplankton taken during the bloom revealed that by Dec 9^th^ *Coscinodiscus* cells were senescing with chloroplastic material coalescing within the cell, compared to earlier images taken on Nov 26^th^ where chloroplastic material was generally intact (Figure 5B). This was 8 days prior to our plaque assay sampling date (Dec 17^th^). As the peak in plaque enumeration did not occur with peaks in total abundance of heterotrophic bacteria during our 13-month sampling window (Figure S7C), we conclude that the substantially higher number of plaques detected in Dec 2020 was due to an increased contribution of bacterial antagonists to total bacterial abundance. As one of the bacteria detected during this bloom was *P. alexandrii* (Figure 2D), we tested whether C. *wailesii* is susceptible to *P. alexandrii*. We isolated a local WEC strain of *C. wailesii* (Methods), Inoculation into co-culture with *P. alexandrii* caused a substantial reduction of growth and Fv/Fm of this strain compared to the no *P. alexandrii* control (Figure 5C). Moreover, cells looked less healthy with empty frustules often observed (Figure 5D). Together, our evidence indicates that antagonistic bacteria showing increased abundance during senescence of a bloom of *C. wailesii* are capable of robust growth inhibitory effects against this diatom.

**Fig. 5.**
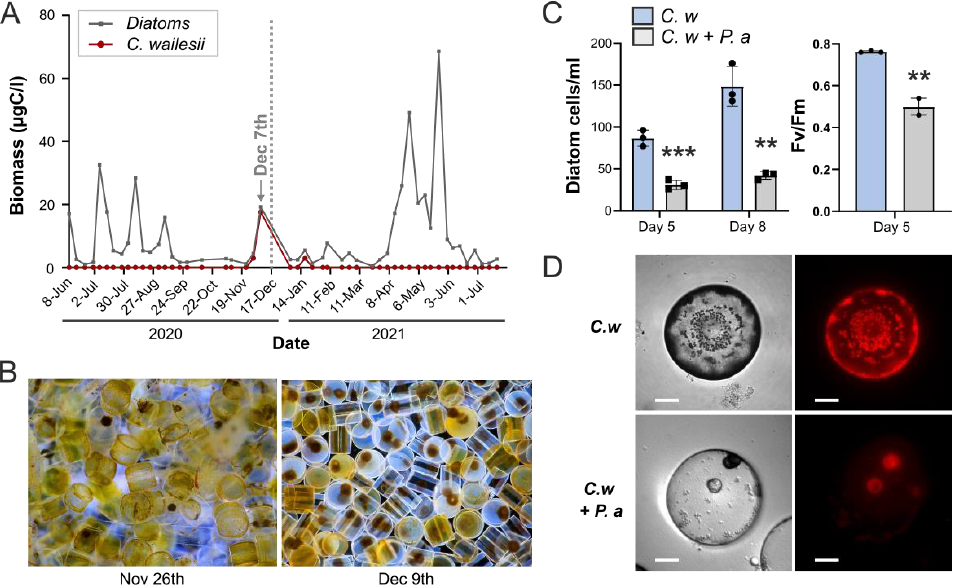
Peaks in plaque enumeration corresponded with senescence of a winter diatom bloom. **A.** Total biomass (µgC/l) of diatoms making up a winter diatom bloom at L4 station in late 2020. The centric diatom *Coscinodiscus wailesii* accounted for the majority of biomass during this bloom. Data was collected by the Western Channel Observatory (WCO) (https://www.westernchannelobservatory.org.uk/). The December plaque assay sampling date (Dec 17^th^, 2020) is indicated with a dashed grey line. The WCO sampling date (Dec 7^th^) just prior to this is also highlighted (grey arrow). The WCO sampling point after this was Jan 4^th^. **B.** Light microscopy images of *Coscinodiscus wailesii* cells sampled from L4 on Nov 26^th^ and Dec 9^th^ (©Richard Kirby). The diameter of *Coscinodiscus* are approximately 350 µm. **C.** Impact *of P. alexandrii* on *C. wailesii* growth and Fv/Fm after 5 and 8 days. On day 8, Fv/Fm values for *C. wailesii* +*P. alexandrii* were undetectable versus 0.55 ± 0.01 in the control. P-values (Student’s t test): **p < 0.01; ***p < 0.001 (*n*=3). **D.** Brightfield images (left) of example *C. wailesii* cells 6 days post inoculation with or without *P. alexandrii*. Epifluorescence images of chlorophyll autofluorescence is also shown (Left). Scale bar: 50 µm.

### Global biogeography of WEC bacterial isolates and their co-occurrence with diatom hosts

To determine the broader biogeography of WEC antagonists, we searched for their distribution in marine ecosystems globally analysing the Ocean Barcode Atlas (40,41). We queried the *Tara* Oceans 16S/18S miTAG metabarcode database with representative 16S rRNA gene sequences of our WEC isolates (Dataset S1). The resemblance of 16S rRNA gene query sequences of WEC isolates, were assessed by compiling maximum likelihood trees for each taxa (Methods; Figure S8-S11). Using this approach, we detected metabarcodes for *T. lohafexi* (or its close relative *T. lucentensis* (42)), *M. adhaerens* and *H. titanicae* (Table S3). While we also identified putative hits for *P. alexandrii* and *M. idriensis*, reads were not detected in the surface water (SRF) or Deep Chlorophyll Maxima (DCM) samples. Additionally, we could not identify close homologues for *M. spongiicola* nor *V. diazotrophicus*. However, we could identify a hit for *M. dokdonensis* (Figure S9). The 16S rRNA gene sequence of which groups together with strong support with *Maribacter* sp. PML-EC2, an algicidal bacterium previously isolated from the WEC that inhibits the growth of *Skeletonema* sp. (CCAP1077/1B; a strain of *S. marinoi* (43)) (39), suggesting that these bacteria are the same species (Figure S9). We therefore examined the distribution of *M. dokdonensis* too. In total, metabarcodes of these four bacterial species were detected in 29 different stations, spanning 7 ocean regions including the S. Pacific, N. Pacific, Mediterranean Sea, Red Sea, Indian Ocean, S. Atlantic and Arctic Oceans (Figure S12; Table S4). *H. titanicae* was the most widespread, identified at 13 *Tara* stations, compared to *T. lohafexi* (12 stations), *M. dokdonensis* (9 stations) and *M. adhaerens* (8 stations). We detected a particularly high incidence of these bacteria in the Mediterranean Sea, with all four represented at station Tara_009, and three of the four (*T. lohafexi*, *M. dokdonensis* and *M. adhaerens*) detected at TARA_007.

To investigate whether we could detect co-occurrence of bacteria with diatom hosts, we also examined the distribution of *T. pseudonana*, *S. marinoi*, and *Chaetoceros* sp. (PLY617) that each exhibit susceptibility to at least one of these bacteria (Figure 4; (39)). Querying 18S rRNA gene sequences for *T. pseudonana* (CCMP1335), we identified two robust hits, with 100% percentage identify (Table S5). *T. pseudonana* was present in the Mediterranean Sea, Indian and Arctic Ocean (examining DCM and SRF). In both the Mediterranean Sea and Arctic Ocean including at the same stations (TARA_22, TARA_168, TARA_173 and TARA_188) we also detected *T. lohafexi* that caused 99.8% inhibition of *T. pseudonana* growth (Figure 6A). These results thus indicate that *T. lohafexi* co-occurs with this diatom host in several different marine ecosystems globally. We also examined co-occurrence patterns of *Skeletonema* sp.. While an 18S rRNA gene sequence is not available for *Skeletonema* sp. CCAP1077/1B that is susceptible *M. dokdonensis* (39), we queried the 18S rRNA gene sequence that was available for another strain of *S. marinoi* (CCMP791), which is very similar if not identical (43). We identified four metabarcodes (% ids ≥98.5%; Table S5) showing a cosmopolitan distribution spanning all ocean regions (Figure 6B) and co-occurring at 6 stations with *M. dokdonensis,* in the Mediterranean Sea and S. Pacific. A maximum likelihood tree grouped three of these metabarcodes with the *S. marinoi* query sequence (Figure S13). However, one metabarcode (18S-miTAG-v2 1614) grouped with *S. menzelii*. We therefore colour coded the *Skeletonema* metabarcode hits: brown (indicating where only *S. marinoi* was detected), yellow (showing where *S. marinoi*-like and *S. menzelii*-like sequences co-occurred) and blue for locations with only *S. menzelii*-like sequences. Using these criteria, our evidence suggests that *M. dokdonensis* co-occurs with *S. marinoi* in the Mediterranean Sea and S. Pacific at two stations (TARA_009 and TARA_82, respectively), and more extensively with *S. menzelii* in these regions (at stations TARA_68, TARA_72, TARA_76, TARA_82 in the S. Atlantic and TARA_007 and TARA_009 in the Mediterranean Sea). Finally, we assessed the distribution of *Chaetoceros* sp., and co-occurrence with *M. adhaerens* and *H. titanicae*. We queried the complete 18S sequence of *C. calcitrans* CCMP1315 (the closest hit to our partial 18S rRNA gene sequence for *Chaetoceros* sp. PLY617) (Dataset S2) (40). This retrieved two metabarcodes, each with % identities of 97.7% (Table S5). While these metabarcodes exhibited a cosmopolitan distribution across multiple *Tara* stations and showed co-occurrence with *M. adhaerens* and *H. titanicae* metabarcodes in several different ocean regions (Figure 6C), the *Chaeotoceros* sp. hits may be a slightly different to PLY617 used in this study (Figure S14).

**Fig. 6.**
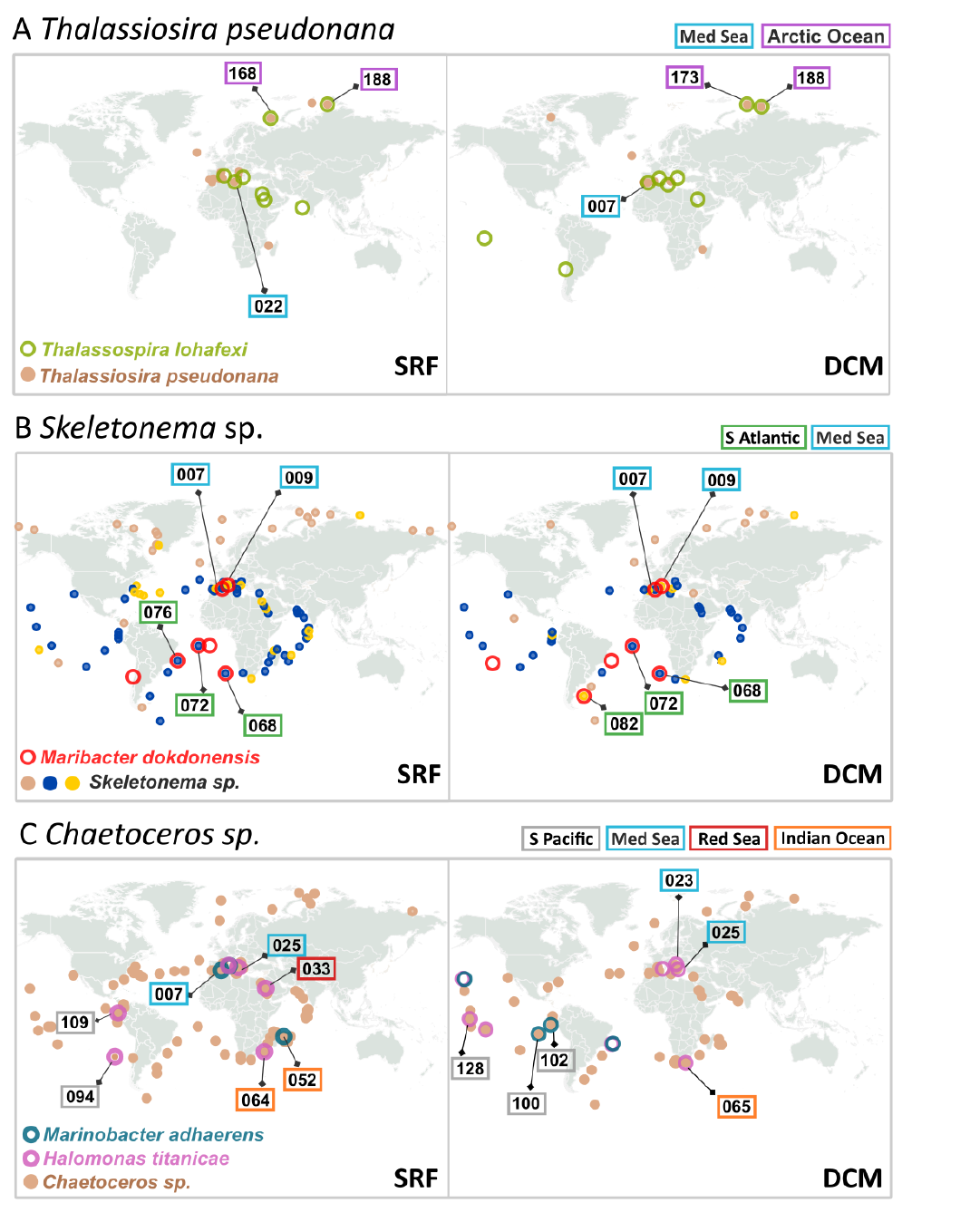
Global biogeography of WEC bacterial isolates with verified antagonistic activity and their co-occurrence with diatom hosts. **A.** Global distributions of *T. lohafexi* and *T. pseudonana* metabarcode hits in the surface (SRF) waters and Deep Chlorophyll Maxima (DCM), as determined through searching the *Tara* Ocean Barcode Atlas (40). TARA stations where co-occurrence of diatom host and bacteria metabarcodes are detected are labelled on the figure. **B.** Distributions of *Maribacter dokdonensis* and *Skeletonema* sp. metabarcode hits are shown, labelled as described in part A. For *Skeletonema* sp., hits are colour coded according to phylogeny: brown indicates where only metabarcodes resembling *S. marinoi* were found, yellow shows locations where both *S. marinoi*-like and *S. menzelii*-like sequences were observed, and blue indicates locations where only *S. menzelii*-like sequences were represented. **C.** Distributions of *Marinobacter adhaerens* and *Chaetoceros* sp. metabarcode hits are shown, labelled as described in part A. D.

In conclusion, we report the widespread distribution of several WEC isolates, and show their cooccurrence with diatom hosts in different ocean regions, for interactions validated in the lab. However, given that bacteria can confer growth inhibitory effect with broad host specificity, it is likely that co-occurrence between bacterial antagonists and susceptible diatoms is more widespread.

## Discussion

Despite accumulating evidence that bacterial pathogens can profoundly impact the growth and physiology of diatoms (8,19), their prevalence and dynamics in natural marine ecosystems is poorly understood. To address this fundamental knowledge gap, we have conducted a 13-month sampling effort to systematically characterise the diversity and seasonal patterns of diatom antagonistic bacteria in a productive coastal ecosystem, where diatoms frequently bloom. We report that multiple and diverse bacterial antagonists co-occur with bloom-forming diatoms in this region. The identification of numerous bacterial species that had no prior reports of antagonistic activity towards diatoms, coupled with our observation that this activity was evident only when antagonists were pre-exposed to diatom organic matter, indicates that bacterial pathogenicity towards diatoms is more prevalent than previously recognised. Moreover, our study captures important evidence of increased detection of bacterial pathogens during senescence of a diatom bloom. We subsequently isolated the major diatom dominating the bloom, *C. wailesii*, and confirmed its susceptibility to *P. alexandrii* that was identified in multiple plaques obtained during this bloom. *C. wailesii* in co-culture with *P. alexandrii* exhibited similar ‘symptoms’ to those observed to in situ *Coscinodiscus* cells visualised just prior to our sampling point that detected increased plaque numbers. Finally, we identified that many of the WEC pathogens identified in this study exhibit geographical ranges extending beyond the WEC, and co-occur with diatom hosts in regions ranging from the Mediterranean Sea to the Arctic Ocean. Our work raises important questions regarding the influences of diatom pathogens on natural diatom populations at the global scale.

Whereas the vast majority of studies of natural diatom populations typically focus on spring in order to capture spring phytoplankton blooms, our study highlights the importance of monitoring the full seasonal cycle. Winter and autumnal diatom blooms are not uncommon, with phytoplankton populations often having a distinct community composition during these seasons (21). In the WEC, the centric diatoms *Coscinodiscus* spp. and *Odontella mobiliensis*, as well as the benthic diatoms *Paralia sulcate* and *Podosira stelligera* dominate winter communities (21). Nanodiatoms including *Minidiscus comicus* and *T. profunda* also peak principally during winter (22). Despite being a key component of WEC phytoplankton communities (22) and global carbon export (44), these tiny diatoms are typically overlooked in routine monitoring due to challenges in their identification. Our observation of peak plaque detection during winter but not spring, suggests that antagonistic bacterial populations may co-occur with, and exert influence on, specific diatom host populations. These important observations provide new opportunities to further dissect the dynamics of antagonistic diatom-bacteria interactions that may be key drivers of diatom blooms and population health in marine ecosystems. This is particularly important in light of evidence that *C. wailesii* (formerly *C. nobilis* Grunow (23)) can have severe detrimental impacts on coastal ecosystems, with extensive mucus production causing clogging of fishing nets in the WEC (45).

The taxonomic composition of our library of WEC antagonistic bacteria was made up predominantly of *Alphaproteobacteria* (44% of species isolated), *followed* by *Gammaproteobacteria* (39%), *Flavobacteria* (11%), and finally members of the *Bacilli* (0.06%). This is in contrast to evidence from a literature of over 100 published papers, which showed that the majority of diatom algicidal bacteria belong to the *Gammaproteobacteria* (42%), compared to just 0.08% representing the *Alphaproteobacteria* (17). However, this review compiled studies of bacteria isolated from a variety of sources. As many reports focused on a single bacterial isolate from their sample/environment, these data likely do not reflect the true make up of antagonist populations in any one marine ecosystem. By sampling frequently and over an annual cycle in the WEC we have gained a greater sampling depth. Our study also addresses a clear bias in sampling efforts towards the study of algicidal bacteria of dinoflagellates (with 116 species reported), compared to those of diatoms (38 bacterial species) (17). Nevertheless, our study may too have been affected by inherent sampling biases. Specific bacterial taxa (e.g. culturable versus unculturable species) are likely more amenable to plaque assay isolation than others. A conspicuous absence of several of the ‘usual suspects’ was also apparent. We did not detect any *Croceibacter* (19) or *Cytophaga*, and only one *Alteromonas* and one *Vibrio* species was isolated, despite these taxa being frequently reported as algicidal (7). It is possible that these bacteria are more representative of oceanic systems (19) or the distinct geographic regions (46,47) from which they were isolated. Certainly, our isolation of *P. alexandrii* on 19 independent occasions from the WEC, despite only one prior report of this species being algicidal (31), suggests it maybe be particularly prevalent in coastal WEC waters.

The predominance of *Alphaproteobacteria* amongst our library of WEC antagonists, challenges current perceptions of the ‘good’ versus ‘bad’ members of the diatom microbiome. With *Alphaproteobacteria* frequently observed to facilitate diatom growth through provision of nutrients or exerting a protective effect (12–15). Similarly, we observed a robust growth inhibitory effect of *M. adhaerens* against *Chaetoceros* sp. PLY617, despite this bacterium more commonly being considered a diatom commensal (48) (albeit a subtle inhibitory effect on *Coscinodiscus radiatus* growth by this bacterium was recently reported (34)). Indeed, this effect was only observed when cells were first grown on medium made up of autoclaved diatoms. By comparison, *M. adhaerens* pre-grown on ½YTSS medium appeared to stimulate diatom growth. This concept of a ‘sliding scale’ from synergistic to antagonistic is not new in marine microbial ecology. Jekyll-and-Hyde dynamics towards coccolithophores have also been observed for *Rhodobacteracae* bacteria *Phaeobacter* (49) and *Sulfitobacter* (50,51). However, our evidence suggests this ‘switch’ towards antagonistic behaviour maybe more common than previously recognised, with the growth inhibitory effects of bacterial species spanning 7 different bacterial orders tested being enhanced by pre-growth on DDM. However, we also observed differences in the dynamics of our system. Whereas *Sulfitobacter* D7 pre-grown on ½YTSS ‘coexisted’ with the coccolithophore *Emiliania huxleyi* for ∼7 days before becoming pathogenic (51), ½YTSS-grown *P. alexandrii* did not cause a culture crash of susceptible diatoms even after 16 days. In the *E. huxleyi*-*Sulfitobacter* system, the signalling molecule algal derived dimethylsulfoniopropionate (DMSP) was identified to be pivotal for mediating the transition to pathogenicity, whereas benzoate hindered this switch (51). Whether bacterial antagonists perceive different cues depending on the algal host remains an open question, particularly as diatoms are typically considered ‘low DMSP producers’ compared to coccolithophores (52). Similarly, the mechanism by which *P. alexandrii* and other WEC isolates confer growth inhibitory effects requires further attention. The algicidal compounds released by *Phaeobacter* species (roseobacticides) (49) are not synthesised by all *Roseobacter* species (53), and algicidal mechanisms are multifaceted (16). The isolation of multiple, diverse bacterial antagonists cooccurring in the WEC offers exciting new opportunities to examine the evolution and ecology of pathogenic strategies amongst bacteria occupying the same niche.

*Alpha*- and *Gammaproteobacteria* (as well as *Flavobacteria*) are often reported to be associated with diatoms and diatom blooms. Indeed, accumulating evidence suggests that representatives of these groups are specialized for decomposition of algal derived organic matter (54), and certain species including *M. adhaerens* physically interact with diatom aggregates (55,56). However, the roles of algicidal bacteria in controlling natural diatom blooms are not well understood. Mesocosm experiments inoculating the algicidal bacterium *K. algicida* into plankton samples taken during a North Sea diatom bloom caused rapid decline of *Chaetoceros* populations (57). However, whether bacterial populations reach sufficient densities in nature to confer such an effect remains an open question. This is particularly pertinent given that density-dependent quorum sensing has been shown to regulate growth inhibitory effects of *Ponticoccus* sp. *PD-2* against the dinoflagellate *Prorocentrum donghaiense* (31). Intriguingly, we observed peaks in plaque detection that co-occurred with senescence of a winter diatom bloom. As total levels of heterotrophic bacteria did not reach peak abundance at the same time, our data suggest enrichment of antagonistic bacteria and/or their activity during this bloom. Further work is now required to determine whether such taxa contribute to diatom bloom demise or are opportunistically scavenging on decaying organic matter as a bloom wanes. Reports of declining diatom populations in the WEC (21) must also be considered in light of our findings that multiple and persistent diatom pathogens inhabit this region.

## Materials and Methods

### Diatom strains and culture conditions

The diatom strains *T. pseudonana* (PLY693; CCMP1335), *T. weissflogii* (PLY541), *Skeletonema sp.* (PLY627), *Chaetoceros sp.* (PLY617) and *P. tricornutum* (PLY100) were obtained from the Marine Biological Association Culture Collection (Plymouth, UK) (Table S1). *C. wailesii* was isolated during this study from the WEC. As *Skeletonema sp.* (PLY627) and *Chaetoceros sp.* (PLY617) were identified only to genus level, 18S rRNA gene sequences were amplified via PCR using the primers Euk1A (CTGGTTGATCCTGCCAG) and EUKB (TGATCCTTCTGCAGGTTCACCTAC), and sequences are given in Dataset S2. All diatoms was cultured in filtered WEC seawater supplemented with f/2 nutrients (including Na_2_SiO and vitamins) (58), at 18 °C (15 °C for *C. wailesii*) under a 16:8-hour light-dark cycle with a light intensity of 30-80 μmol m^2^ s^−1^.

### Isolation of C. wailesii

*C. wailesii* cells were isolated from a natural phytoplankton population on the 17^th^ of October 2022. Seawater was collected with a plankton net (200 µm mesh size) deployed at a depth of 5 m from coastal water in the Plymouth Sound, WEC (50° 20’ 0.093” N, 4°, 08’ 0.915” W). *C. wailesii* cells were isolated with a micropipette prepared on a P-97 micropipette puller with a tip diameter of ∼350 µm (Sutter, Novato, CA, USA). Cells were washed by resuspension into 1 ml droplets of sterile seawater three times prior to being placed into aged filtered sterile seawater with f/2 medium with 100 µM silicate, in 4 ml well plates (∼10 cells per well).

### Antibiotic treatment of diatom cultures and verification of axenity

All diatom strains used for this study were treated with a cocktail of antibiotics. Treatment consisted of three sequential rounds (with the exception of *C. wailesii* that only had one treatment round) of three-day incubations of diatom cells in f/2 medium supplemented with 0.7 mg/ml ampicillin, 0.1 mg/ml streptomycin and 0.5 mg/ml penicillin. Treated diatoms were subsequently sub-cultured into fresh f/2 medium without antibiotics and allowed to reach exponential growth phase. To check the axenity of the cultures, a sample of cells were stained with the nucleic acid–specific stain Hoechst (1 µl/ml), incubated in the dark for 30 min at 20 °C in a glass-bottomed dish and viewed under epifluorescent illumination (excitation 395 nm, emission at 460 nm) using a Leica DMi8 inverted microscope (Leica Microsystems, Milton Keynes, UK) with a 63 × 1.4NA oil immersion objective. Bacteria were clearly visible in nontreated control cultures but were not seen in the antibiotic treated cultures used in this study. Cultures were checked frequently (approximately monthly) via epifluorescence staining and plating on Difco Marine Broth plates through the course of the study. On occasions when contamination was detected cells were re-treated with antibiotics and visually inspected as described above. While antibiotic treatment of *C. wailesii* substantially reduced bacterial load, we did not obtain a completely axenic culture, due to the sensitivity of this strain to antibiotics.

### Isolation of antagonistic bacteria from the Western English Channel via plaque assays

Axenic diatom host species were cultured under standard conditions for 14 days for use in plaque assays. For each assay, 100 ml of diatom cells were concentrated by centrifugation at 3500 rpm for 10 mins and cell pellets were re-suspended in 500 µl sterile f/2 medium. For the bacterial inoculum from the WEC, surface water (5 m) samples were collected monthly (where possible) from Station L4 (50° 15.00′N, 4° 13.02′W, Figure S1) in 20 L carboys. For the June 2020 sample, it was not possible for the research vessel to go to sea for logistical reasons (COVID 19 pandemic), instead samples were taken off the coast of Plymouth (50.363338, - 4.139739). Seawater (1 l) was sequentially filtered through 100 µm, 40 µm, 10 µm pluristrainers to remove larger eukaryotic cells and debris. Finally, remaining cells were collected on a 0.22 µm membrane. The membrane was subsequently washed with 1 ml sterile seawater to produce a concentrated filtrate of bacterial cells.

To prepare plaque assays, 100 µl of concentrated bacterial filtrate (or autoclaved filtered seawater for the control) was inoculated into 500 µl of the concentrated 14-day old axenic diatom culture and left to incubate at room temperature for 30 mins to 1 hr. The diatom-bacteria co-culture was added to 4.4 ml of 0.4 % molten (55°C) f/2 agar (f/2 medium supplemented with 0.4 g/l agarose) and mixed thoroughly by inversion. Agar was poured immediately onto a 1% f/2 agar plate, incubated at 18 °C under a 16:8 hour light-dark cycle with a light intensity of 50-80 μmol m^−2^ s^−1^ and observed for several weeks for plaque formation.

For the plaque-to-plaque assay experiments described in Table S2, upon plaque formation on the original plaque assay plate, single plaques were picked with a sterile loop, and immediately used to create a new inoculum by placing into 500 µl sterile seawater, 100 µl of which was inoculated into a fresh plaque assay with each of the diatom hosts. The number of plaques was then quantified after a minimum of four weeks.

### Identification of bacterial isolates from plaque assays

Plaques picked from successful plaque assays were streaked onto 1% agar plates made up of ½YTSS medium (containing 2 g/l of yeast extract, 1.25 g/l of tryptone, and 20 g/l of sea salts (Sigma). Pure cultures were obtained by streaking single colonies a minimum of four times. In instances where two colony types were apparent from a plaque streak out (P45, P47, P46, P49 and P29) (Figure S2), both colonies were processed and sequenced. Bacterial 16S rRNA gene was amplified via a GoTaq (Promega, USA) colony PCR using primers 27f (5′-AGAGTTTGATCMTGGCTCAG-3′) and 1492r (5′-TACGGYTACCTTGTTACGACTT-3′) (59,60), and the following PCR cycle conditions: initial denaturation at 95 °C for five mins, followed by 32 cycles of i) denaturation at 95 °C for 30 secs, ii) annealing at 50.1 °C for 30 secs and iii) extension at 72 °C for 90 secs. A final extension at 72 °C was then carried out for five mins. 16S rRNA PCR products were purified using a QIAquick PCR Purification kit (Qiagen, UK) and sequenced via Sanger sequencing by Source Bioscience (Cambridge, UK). Retrieved sequences were trimmed and aligned using MEGA11 (61) and closest hits were identified using the NCBI Basic Local Alignment Search Tool (BLAST) nucleotide database (62). The phylogenetic classification of hits was then validated via construction of Maximum Likelihood trees with type strains from the relevant genera obtained via the ribosomal Database Project (see *Maximum Likelihood tree* methods section).

### Physiological growth assays co-culturing diatoms with DDM- and ½YTSS-grown WEC bacteria

Prior to inoculation of co-culture experiments, bacteria were pre-grown for four days at 18 °C on ½YTSS or DDM agar plates. DDM plates were prepared by autoclaving 14-day old axenic diatom cultures with 1% agarose, and used the same day. For all experiments, DDM was prepared from the diatom host the bacterium was to be inoculated into co-culture with (i.e. *T. pseudonana* DDM for *T. pseudonana-P. alexandrii* co-cultures). Except for *C. wailesii-P. alexandrii* experiments in which *P. alexandrii* was pre-grown on *T. pseudonana* DDM plates. Bacteria pre-grown on ½YTSS or DDM plates were collected with a sterile loop and re-suspended in 1 ml (0.2 µm) filtered seawater without nutrients. This bacterial suspension was inoculated into fresh f/2 medium to a final optical density (OD)600 of 0.05, and host diatoms were inoculated to a concentration of 30,000 cells/ml (with the exception of *C. wailesii* where the starting density was 8 cells/ml). Diatom cell density was measured during exponential phase of diatom growth using a haemocytometer (or Sedgewick Rafter cell for *C. wailesii*), and maximum quantum efficiency of photosystem II (Fv/Fm) monitored using a PSI AquaPen AP100 (Photon Systems Instruments, Czech Republic). For experiments determining viability of *T. pseudonana* cells in co-culture with *P. alexandrii*, bacterial cells was pre-grown on liquid *T. pseudonana* DDM + 0.1% glucose prior to being inoculated into co-culture. Diatom cell counts were taken using a Luna Automated Cell counter (Logos Biosystems), using SYTOX Green stain NucGreen™ Dead 488 (ThermoFisher Scientific).

### Bioinformatics assessment of the global biogeography of WEC bacterial antagonists and diatom co-occurrence

16S rRNA gene query sequences of WEC bacterial antagonists with verified growth inhibitory effects against diatoms, including *T. lohafexi*, *P. alexandrii*, *M. dokdonensis*, *H. titanicae*, *M. adhaerens*, *M. idriensis*, *M. spongiicola* and *V. diazotrophicus* (Dataset S1) were searched against the *Tara* Oceans 16S/18S miTAG metabarcode database via the Ocean Barcode Atlas portal (40) using the ‘vsearch’ method. Top hits detected in SRF or DCM samples were further scrutinised to assess their taxonomic classification by constructing maximum likelihood trees with query and metabarcode hit sequences as well as sequences of Type strains of closely related bacteria, obtained via the ribosomal Database Project (see *Maximum Likelihood tree construction* methods section below). Distributions of bacterial metabarcodes or ‘miTAGs’ clustering with query sequences of our WEC bacteria were subsequently plotted, pooling data from the available 0.22-1.6 µm and 0.22-3 µm size fractions, including for *T. lohafexi*, *M. dokdonensis*, *H. titanicae* and *M. adhaerens*.

To examine distribution patterns of diatom hosts, 18S rRNA gene query sequences for *T. pseudonana* CCMP1335, *S. marinoi* (CCMP791) and *Chaetoceros calcitrans* (CCMP1315) (NCBI identifiers: AY485452.1, AB948146.1 and KY852256.1, respectively) were searched against the in the *Tara* Oceans OTU 18S V9 database (40). Metabarcodes from samples of different size fractions (including 0.8-5 µm, 0.8-20 µm, 5-20 µm, 20-180 µm, >0.8 µm and >3 µm) were pooled, and their distribution examined. For *T. pseudonana,* the metabarcodes retrieved exhibited a 100% percentage identity with the query sequence and so no further phylogenetic analysis was performed. However, for the *S. marinoi* and *Chaetoceros* sp. searches, retrieved hits exhibited a percentage identity below 100% and so all metabarcodes displayed in Figure 6B-C were subjected to further examination to assess their phylogenetic affiliation via construction of Maximum Likelihood trees made up of the query sequence, metabarcode hits and a selection of taxa from each respective genus, as described below.

### Maximum Likelihood tree construction

Multiple sequence alignments of sequences were generated via Multiple Alignment using Fast Fourier Transform (MAFFT) tool (63). After manual refinement, Maximum Likelihood trees were generated using MEGA11 (61), using the General Time Reversable model with 1000 bootstraps.

### Assessment of diatom diversity and abundance at L4 station

To determine diatom abundance and diversity at L4 station seawater samples taken weekly (wherever possible) from a depth of 10 m, were fixed with acid Lugol’s iodine solution (2% final concentration) and examined by light microscopy via the Utermöhl counting technique (21). All metadata are freely available from the WCO (https://www.westernchannelobservatory.org.uk/). Plankton samples for the *Coscinodiscus* images were collected at the location 50.281936, −4.076956, near Plymouth, UK, by horizontal tow at a depth of ∼ 5 m using a 280 µm mesh plankton net with a 30 cm diameter mouth.

## Acknowledgements

We acknowledge support from the NERC Independent Research Fellowship grant NE/R015449/2 (K.E.H.) and PhD studentship from the NERC ARIES doctoral training program (L.B.). C.E.W. was supported by funding from the NERC National Capability Long-term Single Centre Science Programme, Climate Linked Atlantic Sector Science, grant number NE/R015953/1. We also thank crew of the research vessel Sepia (MBA, Plymouth, UK) for their collection of samples used during this study.

