## Supplementary Information for "Cryptic bacterial pathogens of diatoms peak during senescence of a winter diatom bloom"

Supplementary Figures and Tables

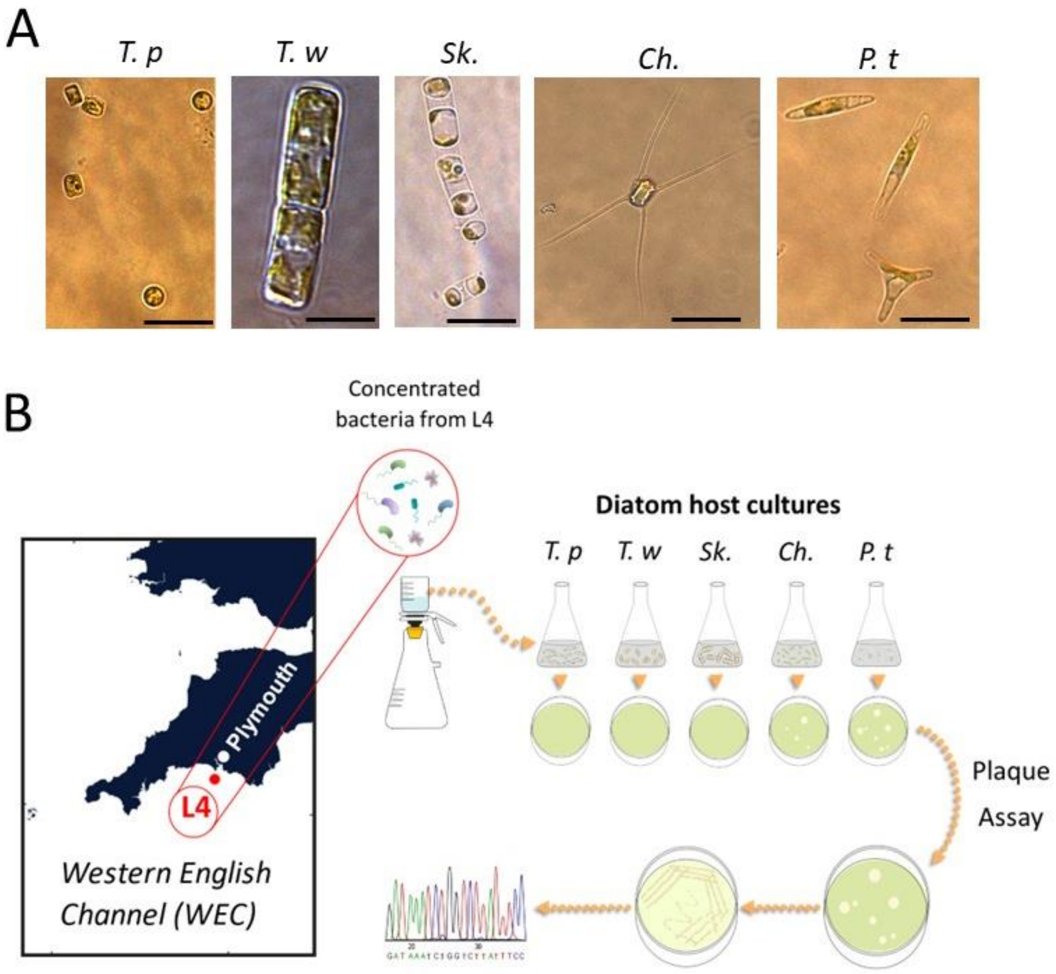


**Figure S1. Schematic diagram of the environmental sampling pipeline used to systematically isolate diatom antagonistic bacteria from the Western English Channel (WEC). A.** Differential interference contrast (DIC) images of diatom species used in this study for plaque assays, including: *Thalassiosira pseudonana* (*T. p*), *Thalassiosira weissflogii* (*T. w*), *Skeletonema* sp. PLY627 (*Sk.*), *Chaetoceros* sp. PLY617 (*Ch.*), as well as *Phaeodactylum tricornutum* (*P. t*). Scale bar: 10 µm. **B.** Plaque assays inoculating diatom hosts with the bacterial fraction of seawater (seawater sampled from L4 station (WEC)), were conducted approximately monthly between June 2020 and July 2021. To identify bacteria from plaques, material from plaques was picked and streaked sequentially 4 times onto 1% ½YTSS agar plates to obtain single colonies and the 16S rRNA gene amplified and sequenced.

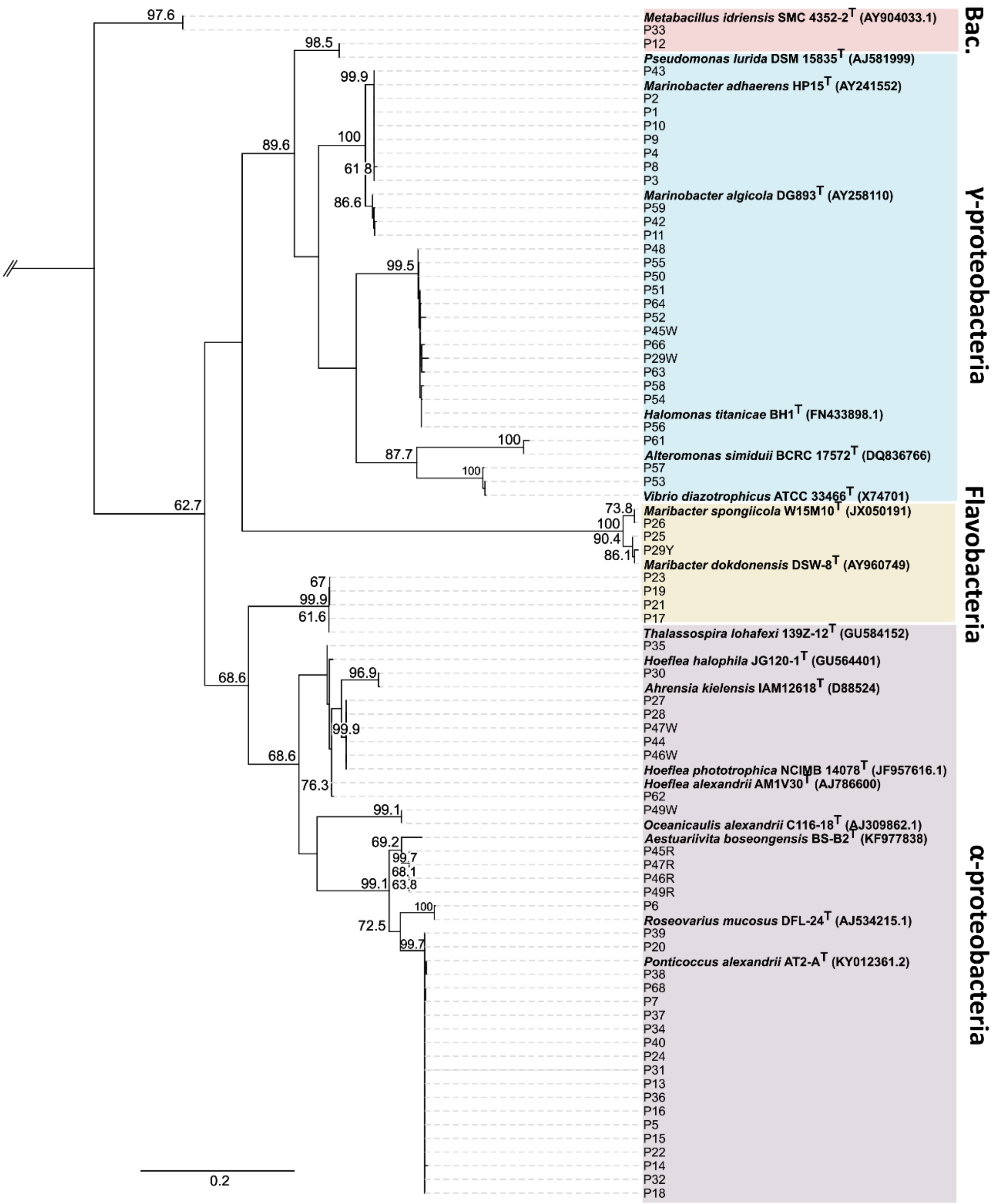


**Figure S2. Maximum likelihood phylogenetic tree of 16S rRNA gene sequences from bacteria isolated from 65 plaques identified during this study.** Sequences obtained from individual plaques are labelled. For five plaques, two colony types were observed, which were sequenced and included in the tree labelled with 'R' (red), 'W' (white), 'Y' (yellow). Reference sequences from Type strains of closely related taxa (defined as described in the methods) are also included (bold). Support values (%) from 1000 bootstrap replicates are indicated below each branch; only values above 60% are given. The 'P' number represents the plaque ID from which the bacterial sequence was obtained.

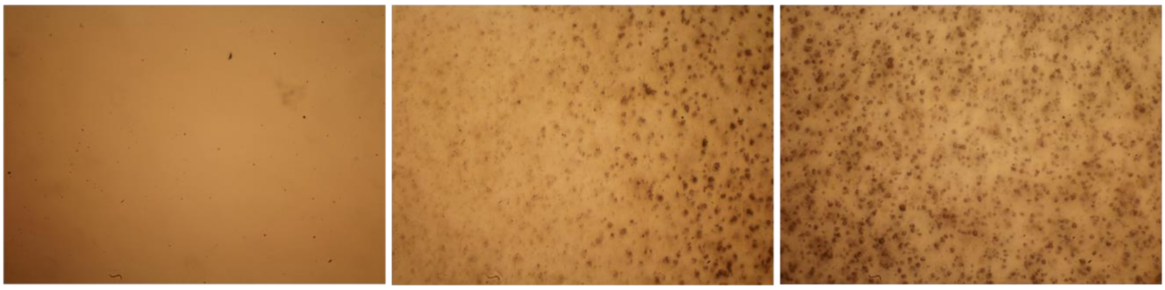


Figure S3. Light microscope image of an example plaque on a *Chaetoceros* plaque assay plate. The image shows the centre of the plaque (left), the plaque boundary (middle) and the diatom lawn outside of the plaque (right).

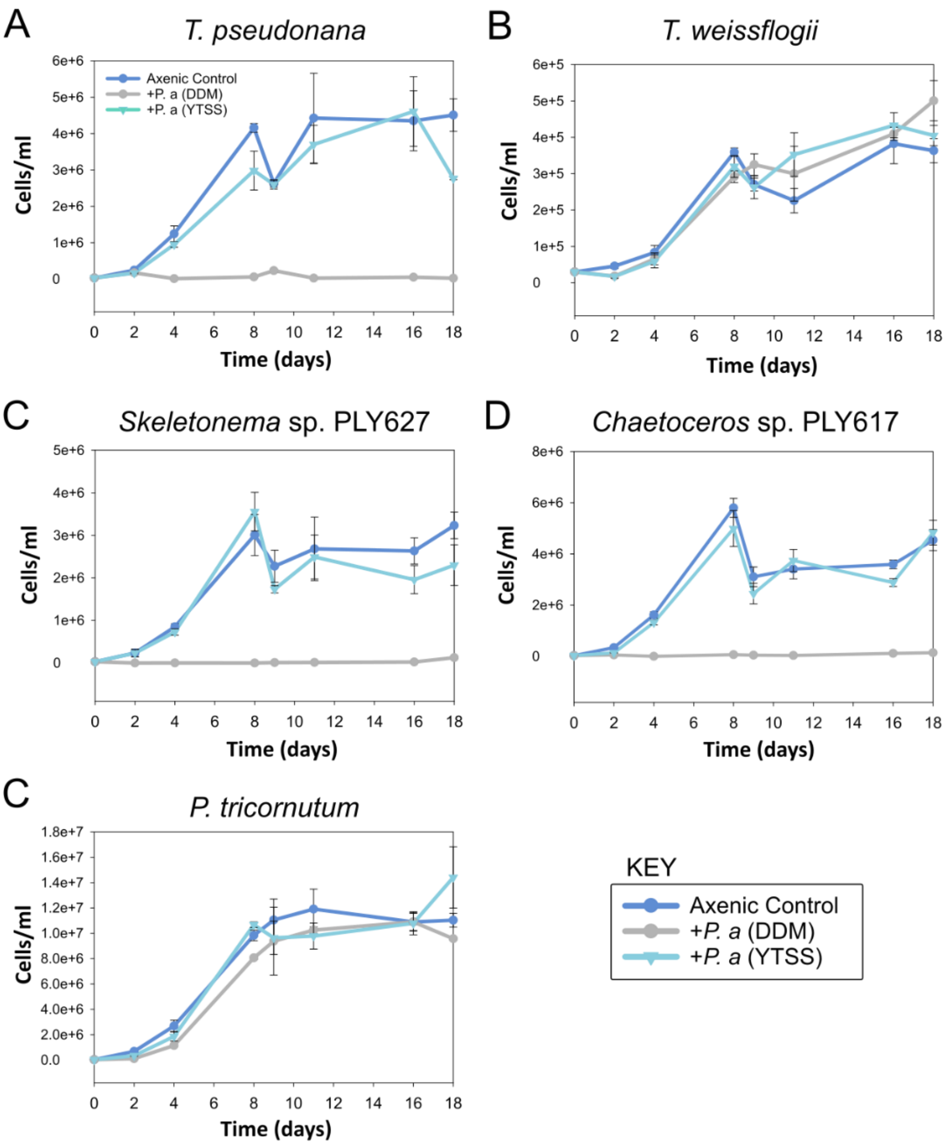


**Fig. S4. Growth over time of five axenic diatom host species in co-culture with *Ponticoccus alexandrii*.** A growth curve of *T. pseudonana* (A), *T. weissflogii* (B), *Skeletonema* sp. PLY627 (C), *Chaetoceros* sp. PLY617 (D) and *P. tricornutum* (E) inoculated with *P. alexandrii* (*P. a*) pre-grown for four days on either DDM or ½YTSS medium. Error bars indicate ± S. E. M for n= 3 replicates.

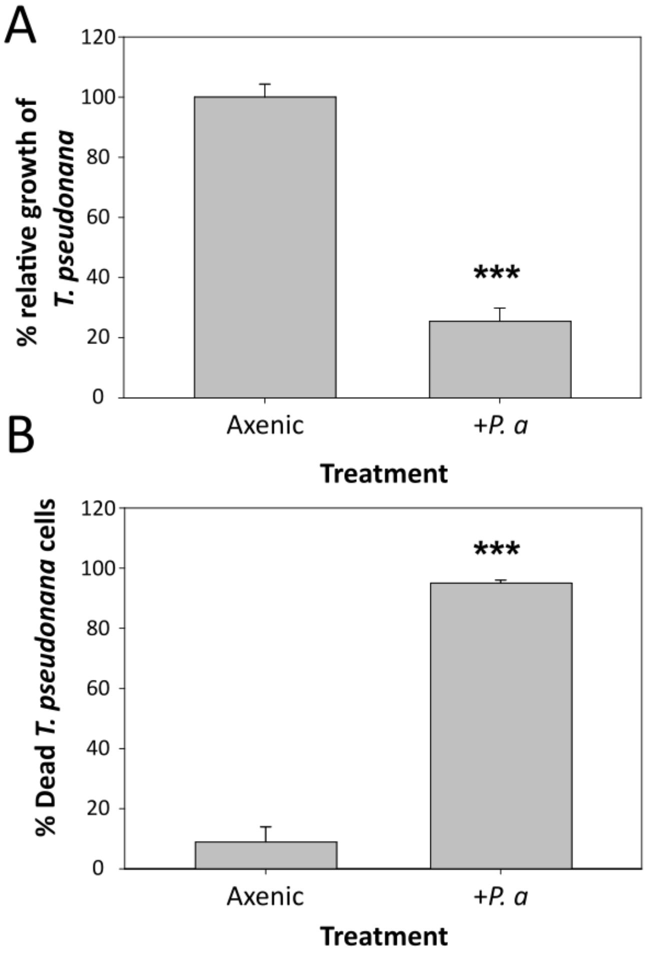


Fig. S5. *P. alexandrii* is algicidal towards *T. pseudonana*. A. Percentage growth (after 4 days) of *T. pseudonana* (relative to the axenic control) inoculated in co-culture with *P. alexandrii* pre-grown on liquid DDM (supplemented with + 0.1% glucose). Error bars indicate ± S. E. M for n=4. P-values (two-tailed t-test): ***$p < 0.001$ (*n*=4). B. Percentage of dead *T. pseudonana* cells for cultures described in (A). Error bars indicate ± S. E. M for n= 4 replicates. P-values (two-tailed t-test): ***$p < 0.001$ (*n*=4).

.

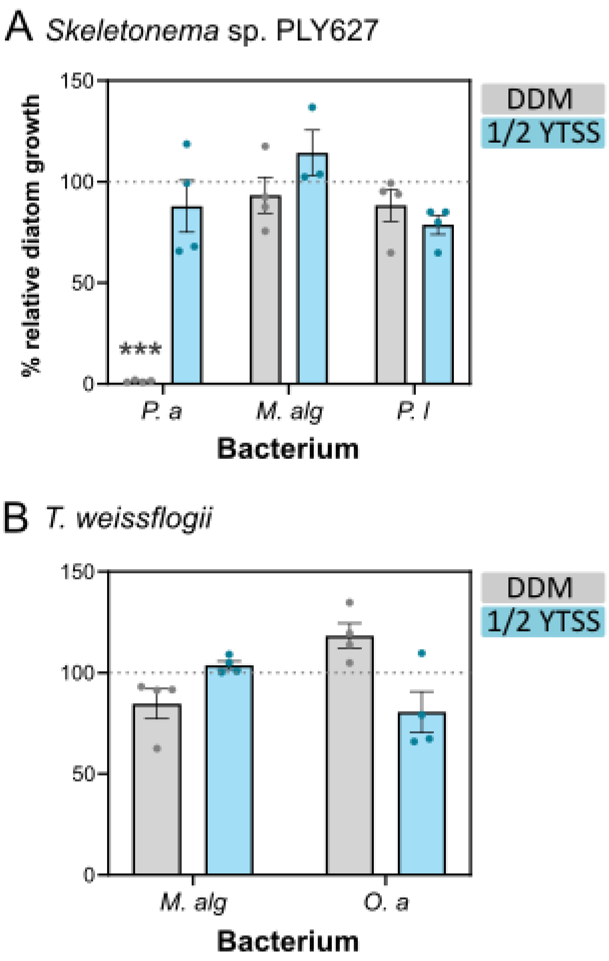


**Fig S6. Growth of the diatoms *Skeletonema* sp. PLY627 and *T. weissflogii* in co-culture with different WEC plaque-forming bacteria.** Percentage relative growth of the diatoms *Skeletonema* sp. PLY627 (**A**) and *T. weissflogii* (**B**) in the presence of plaque-forming WEC bacteria relative to the axenic control. Bacterial isolates tested included: *Ponticoccus alexandrii* (*P. a*), *Marinobacter algicola* (*M. alg*), *Pseudomonas lurida* (*P. l*), and *Oceanicaulis alexandrii* (*O. a*). The effect on the growth of diatom host on which the bacterium was originally isolated via plaque assay was tested. Error bars indicate ± S. E. M for n=4 or 3; P-values (one-way ANOVA comparing co-culture treatments to the axenic control): $P < 0.05$; $^{**}p < 0.01$; $^{***}p < 0.001$ (*n*=3 to 4). Growth was quantified by measuring diatom cell density (cells/ml) following inoculation into fresh f/2 media +Si (to a final cell density of 30,000 cells/ml) with or without WEC bacteria (whereby bacteria in '+bacteria' treatments were inoculated to a final optical density ($OD_{600}$) of 0.05.

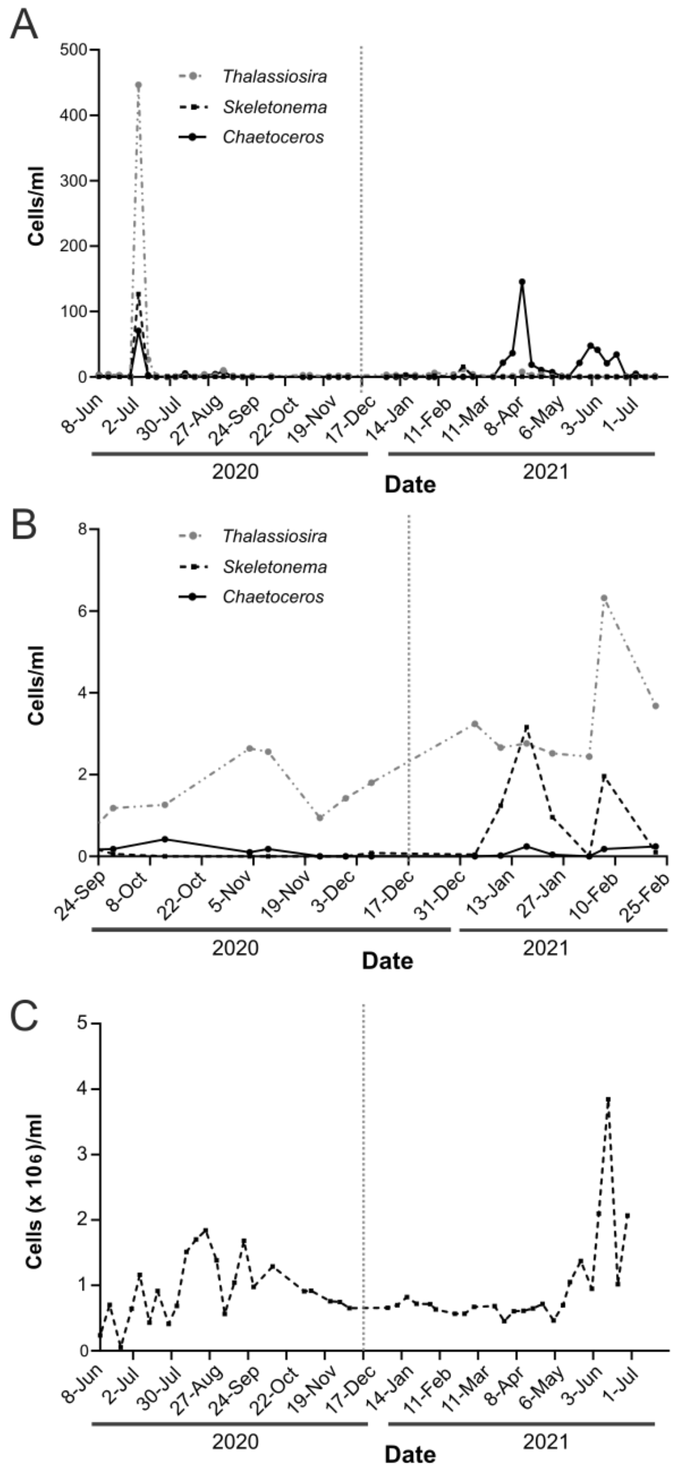


Figure S7. Abundance of diatoms of the genera *Skeletonema*, *Thalassiosira* and *Chaetoceros* as well as bacteria at L4 station from May 2020 to June 2021. Abundance (cell/ml) of diatoms belonging to the genera *Skeletonema*, *Thalassiosira* and *Chaetoceros* between June 2020 and July 2021 (**A**). A more narrow time window is also shown, between the 24$^{th}$ of Sept and 25$^{th}$ of Feb (**B**). Abundance of heterotrophic bacterioplankton between June 2020 and July 2021 is shown in (**C**). Data was collected as part of the Western Channel Observatory (WCO). The December plaque assay sampling date (Dec 17$^{th}$, 2020) is indicated with a dashed grey line.

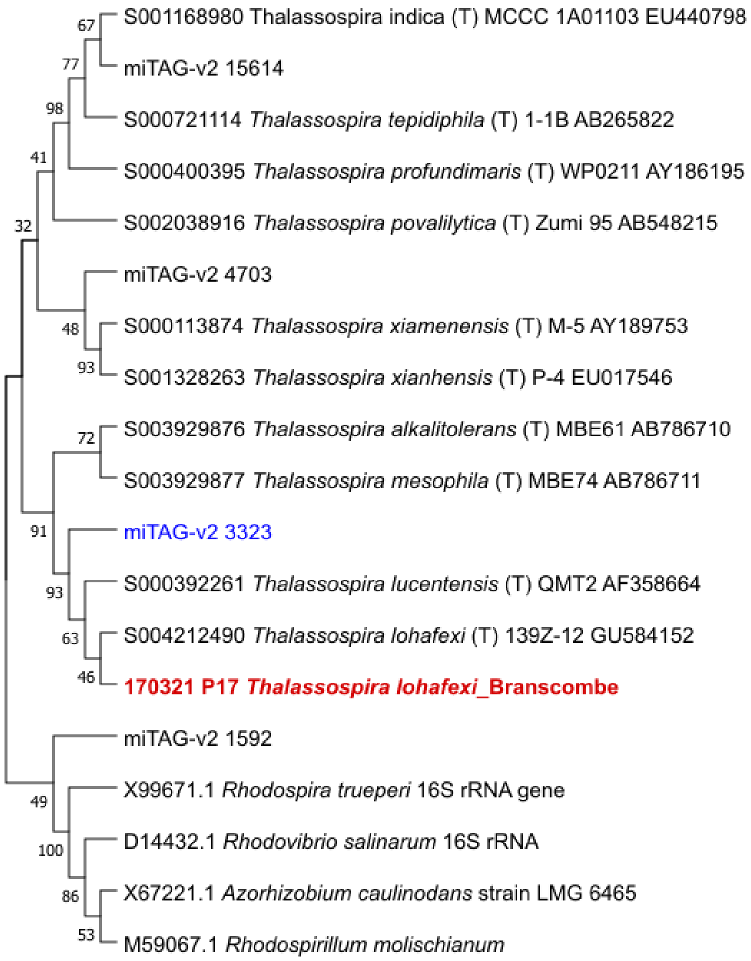


**Figure S8. Maximum likelihood phylogenetic tree of 16S rDNA sequences of *Thalassospira* species.** The tree includes the sequence of the WEC *T. lohafexi* species (obtained from plaque no. 17) isolated in this study (red, bold), in addition to the type strains *T. lohafexi* 139Z-12 and other species of the *Thalassospira* genus (1) (black, bold), as well as sequences resembling *T. lohafexi* identified in the *Tara* 16S/18S rDNA miTAG metabarcode database (2,3) via the Ocean Barcode Atlas (4) (blue). Support values (%) from 1000 bootstrap replicates are indicated below each branch. *Azorhizobium caulinodans strain LMG 6465* was used as an outgroup.

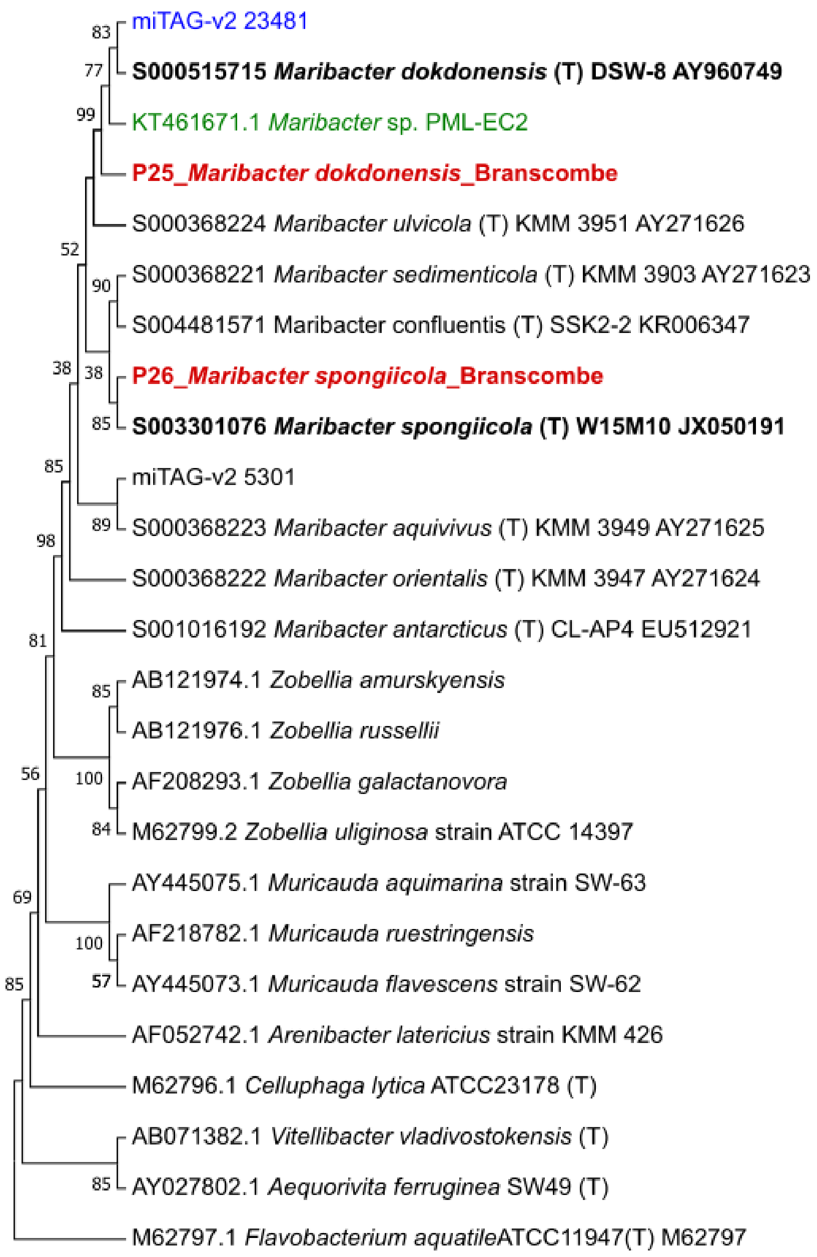


**Figure S9. Maximum likelihood phylogenetic tree of 16S rDNA sequences of *Maribacter* species.** The tree includes sequences of the WEC *Maribacter* species *M. spongiicola* and *M. dokdonensis* (obtained from plaque no. 26 and 25, respectively) isolated in this study (red, bold), in addition to the type strains for *M. spongiicola* W15M10 (5) and *M. dokdonensis* DSW-8 (6) and other species of the *Maribacter* genus (black, bold), as well as sequences resembling *M. spongiicola* and *M. dokdonensis* identified in the *Tara* 16S/18S rDNA miTAG metabarcode database (2,3) via the Ocean Barcode Atlas Atlas (4) (blue). Additionally, the sequence for *Maribacter* sp. PML-EC2 (green), an algicidal bacterium previously isolated from the WEC is also included (7). Support values (%) from 1000 bootstrap replicates are indicated below each branch. *Flavobacterium aquatile* ATCC11947 type strain was used as an outgroup.

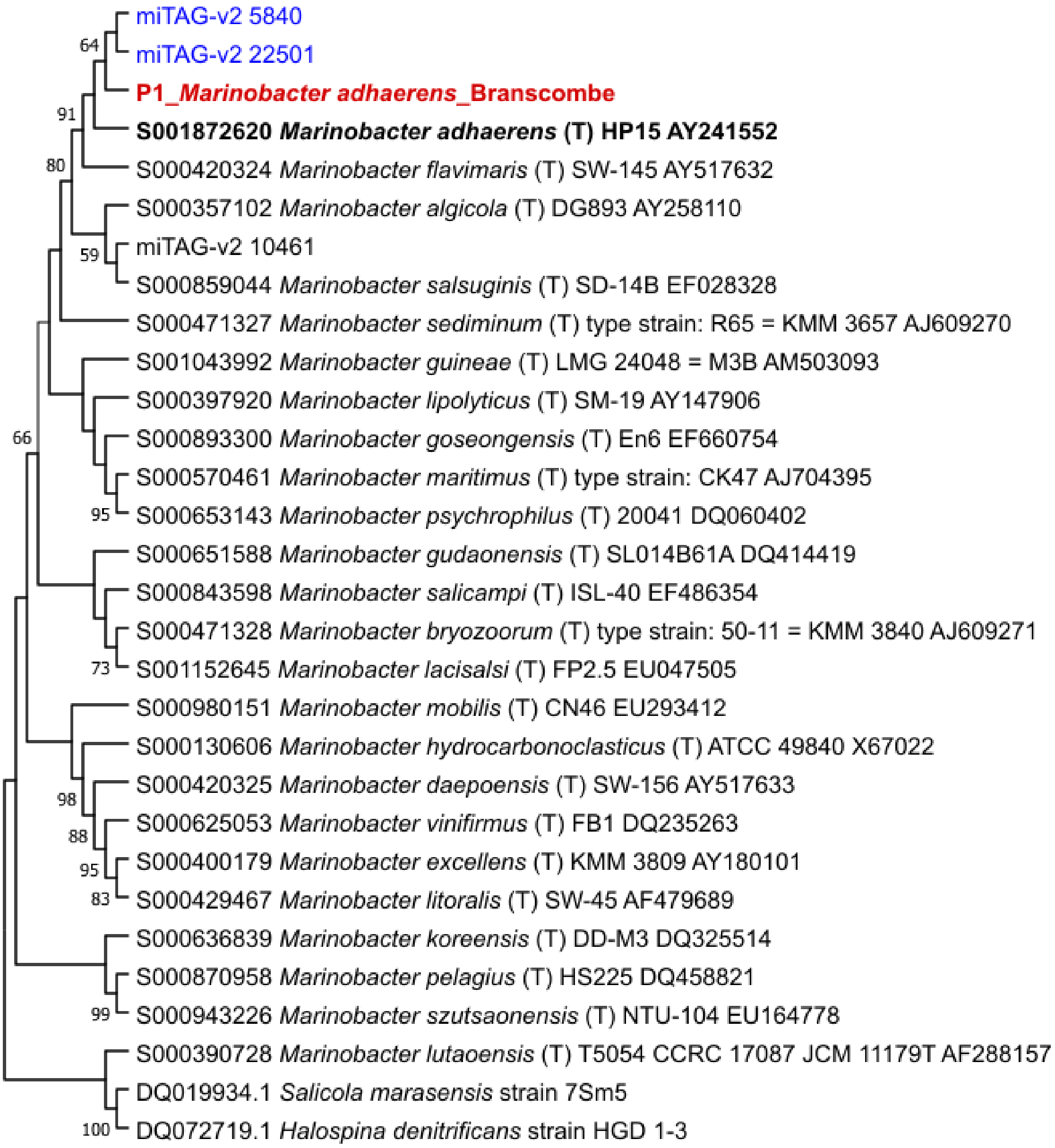


**Figure S10. Maximum likelihood phylogenetic tree of 16S rDNA sequences of *Marinobacter* species.** The tree includes the sequence of the WEC *M. adhaerens* species (obtained from plaque no. 1) isolated in this study (red, bold), in addition to the type strains for *M. adhaerens* HP15 and other species of the *Marinobacter* genus (8) (black, bold), as well as sequences resembling *M. adhaerens* identified in the *Tara* 16S/18S rDNA miTAG metabarcode database (2,3) via the Ocean Barcode Atlas (4) (blue). Support values (%) from 1000 bootstrap replicates are indicated below each branch; only values above 50% are given. *Halospina denitrificans* strain HGD 1-3 was used as an outgroup.

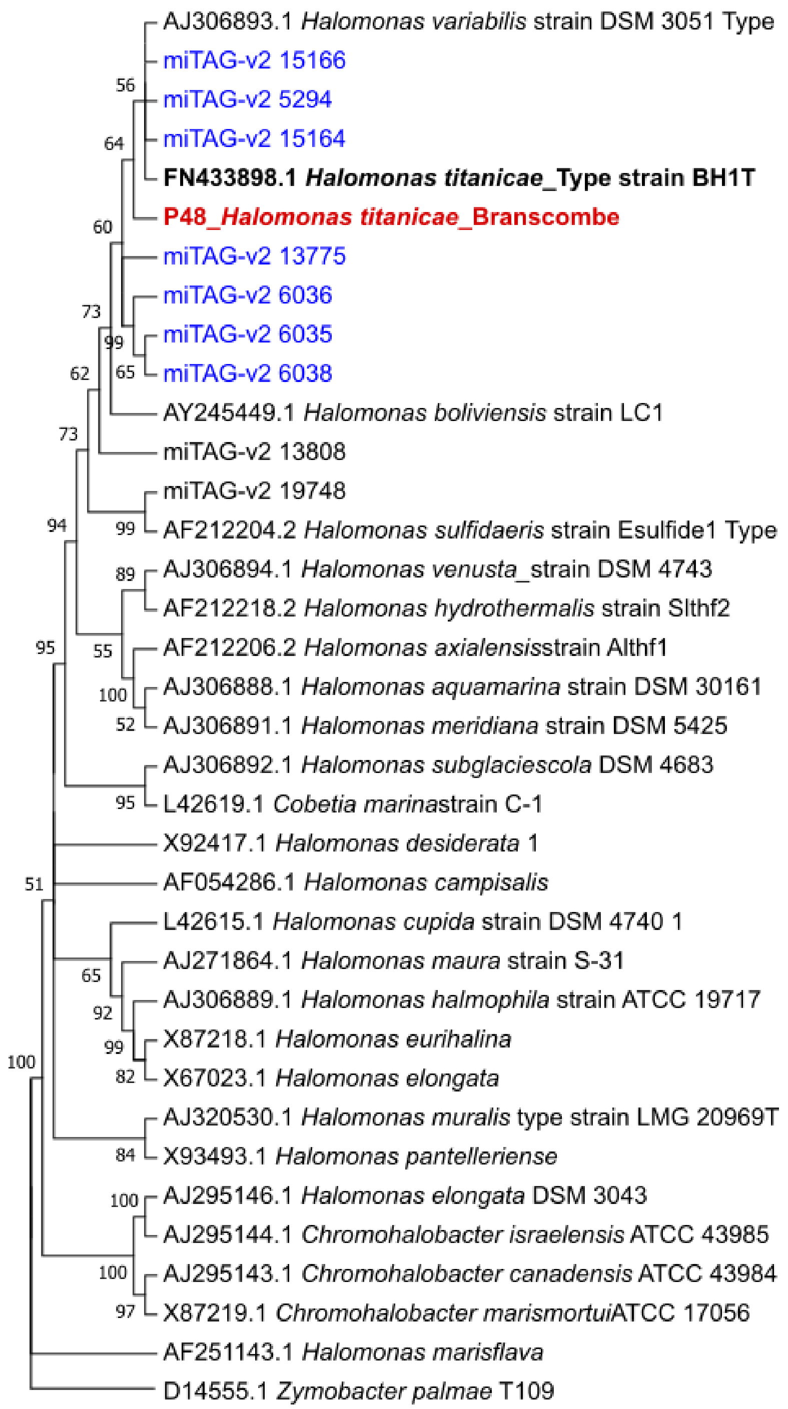


Figure S11. Maximum likelihood phylogenetic tree of 16S rDNA sequences of *Halomonas* species. The tree includes the sequence of the WEC *H. titanicae* species (obtained from plaque no. 48) isolated in this study (red, bold), in addition to the type strains for *H. titanicae* strain BH1T and other species of the (9) (black, bold), as well as sequences resembling *H. titanicae* identified in the *Tara* 16S/18S rDNA miTAG metabarcode database (2,3) via the Ocean Barcode Atlas (4) (blue). Support values (%) from 1000 bootstrap replicates are indicated above each branch; only values above 50% are given. *Zymobacter palmae* D1O9 type strain was used as an outgroup.

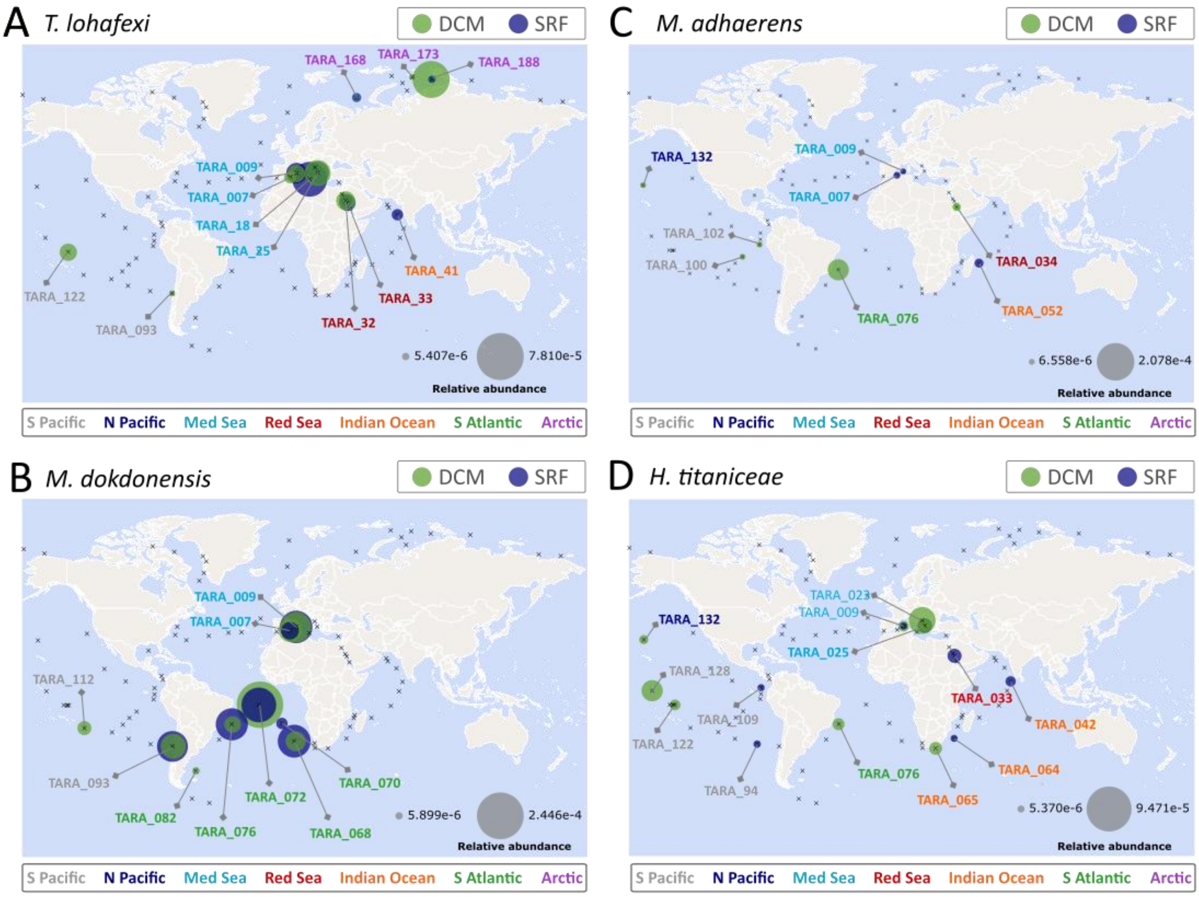


**Figure S12. Global distribution and abundance of metabarcodes resembling WEC antagonists in the *Tara* Oceans 16S/18S rDNA miTAG metabarcode database.** Identifiers for the metabarcode/s plotted for each species are listed in Table S3. Reads were pooled from the 0.22-1.6 µm and 0.22-3 µm fractions. The relative abundance of each metabarcode was estimated via the Ocean Barcode Atlas portal Atlas (4), by dividing the number of barcode counts by the total number of metabarodes in the corresponding sample. The two grey circles of the map scales represent the maximum and minimum abundance related to the barcode numerical values. Nb. No metabarcodes were detected in the N. Atlantic Ocean.

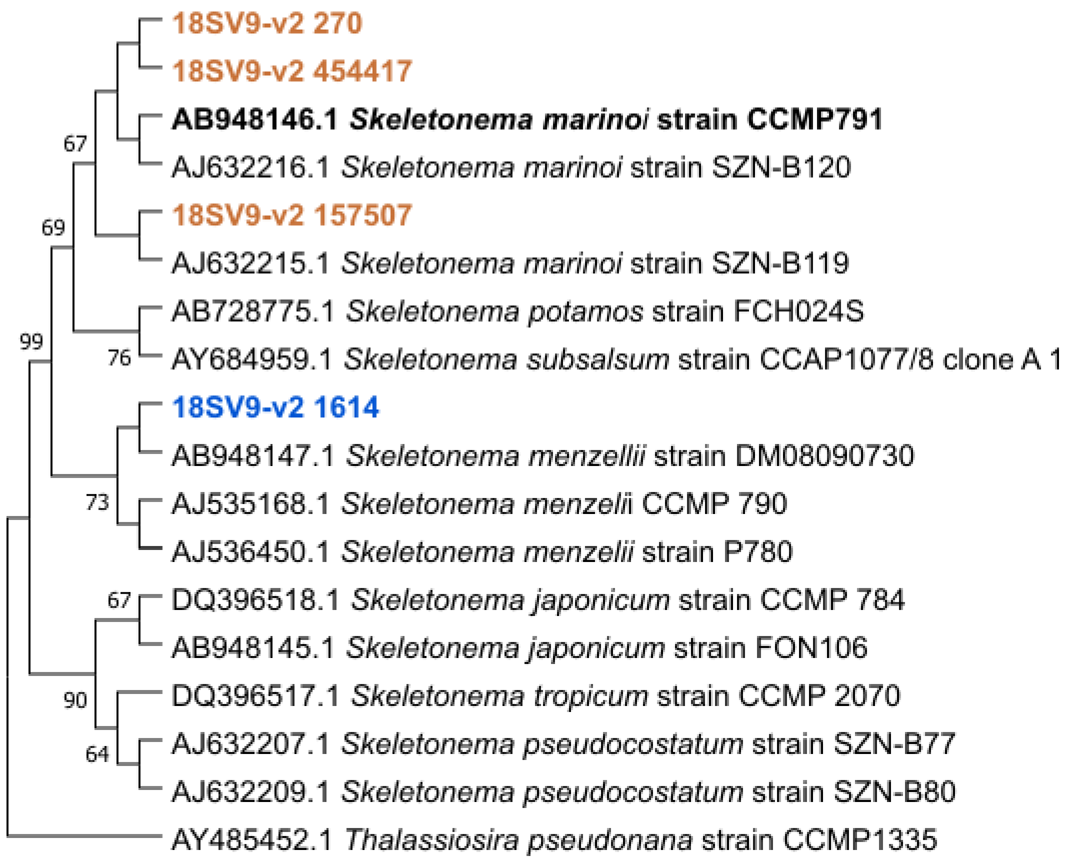


**Figure S13. Maximum likelihood phylogenetic tree of 18S V9 rDNA sequences of *Skeletonema* species.** The tree includes sequences of the *Skeletonema marinoi* strain CCMP791 shown previously to be closely related to *S. marinoi* CCAP1077/1B that is susceptible *M. dokdonensis* (7). Environmental sequences identified by searching the *Tara* Oceans OTU 18S V9 database via the Ocean Barcode Atlas (4) are also included (coloured brown and blue). Support values (%) from 1000 bootstrap replicates are indicated below each branch; only values above 50% are given. *Thalassiosira pseudonana* CCMP1335 was used as an outgroup.

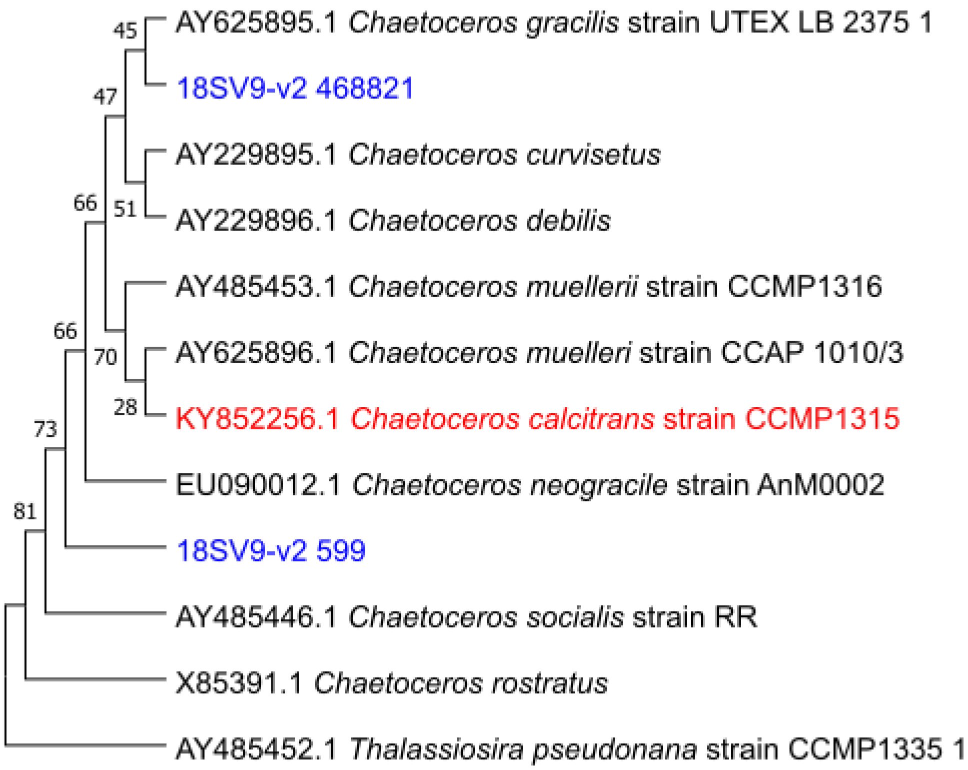


**Figure S14. Maximum likelihood phylogenetic tree of 18S V9 rDNA sequences of *Chaetoceros* sp.** The tree has been constructed from a sample of *Chaetoceros* species including *C. calcitrans* strain CCMP1315 (the closest hit to our partial 18S rDNA sequence for *Chaetoceros* sp. PLY617 that excludes the V9 region), as well as environmental sequences identified by searching the *Tara* Oceans OTU 18S V9 database via the Ocean Barcode Atlas (4) (blue). For both trees, support values (%) from 1000 bootstrap replicates are indicated above each branch, and *Thalassiosira pseudonana* CCMP1335 was used as an outgroup.

**Table S1. Diatom taxa examined in this study.** Strain details as well as isolation location are given.

| Diatom species | Location of isolation | Latitude and longitude |
|---|---|---|
| *Thalassiosira pseudonana* (PLY693; CCMP1335) | Moriches Bay, Forge River, Long Island, New York USA | 40.7560N 72.8200W |
| *Thalassiosira weissflogii* (PLY541) | Gorleston -on Sea, Norfolk, England | N/A |
| *Skeletonema* sp. PLY627 | L4 Station, Western English Channel | 50°15'N 4°13W |
| *Chaetoceros* sp. PLY617 | L4 Station, Western English Channel | 50°15'N 4°13W |
| *Phaeodactylym tricornutum* (PLY100) | English Channel, Off Plymouth | N/A |
| *Coscinodiscus wailesii* | Western English Channel | 50° 20’ 0.093” N, 4°, 08’ 0.915” W |

**Table S2. Re-inoculation of plaques picked from original environmental plaque assays enables propagation of plaques.** Number of plaques formed on each diatom host lawn following plaque assay re-inoculation with plaques identified to contain *P. alexandrii*, *M. adhaerens,* and *M. algicola*, obtained from the original environmental plaque assays. For *M. algicola*, plaques were inoculated back into plaque assays with *T. pseudonana* only.

| **Bacterium** | **Number of plaques per diatom host species** | | | | |
|---|---|---|---|---|---|
| | *T. pseudonana* | *T. weissflogii* | *Skeletonema* sp. (PLY627) | *Chaetoceros* sp. PLY617 | *P. tricornutum* |
| *P. alexandrii* | 38 | 0 | >100 | 47 | 0 |
| *M. adhaerens* | >100 | 0 | 0 | >100 | 0 |
| *M. algicola* | 60 | - | - | - | - |

**Table S3. List of metabarcodes identified in the TARA Ocean Barcode Atlas resembling 16S rDNA sequences of WEC bacterial isolates with verified antagonistic activity against diatoms.** Details are provided on the diatom host/s each bacterium was originally isolated on via plaque assay, as well as the diatom species that each bacterium has been shown in liquid culture to confer a growth inhibitory effect. The *Tara* Ocean barcode Identified for each hit, as well as details of % identity to the query sequence, and no. of reads detected are also given.

| *WEC bacterial isolate* | *Diatom plaque assay species* | *Verified growth inhibitory effect against in liquid culture* | Reference | *TARA Ocean Barcode Id.* | *% identity* | *No. of reads* |
|---|---|---|---|---|---|---|
| *Thalassospira lohafexi* | *T. p, Sk sp. and T. w* | *T. pseudonana (CCMP1335)* | *This study* | *miTAG-v2_3323* | *99.8* | *691* |
| *Ponticoccus alexandrii* | *T. p, Sk sp., Ch. sp. and T. w* | *T. pseudonana (CCMP1335)* | *This study* | *miTAG-v2_22646* | *99.6* | *1* |
| *Maribacter dokdonensis* | *P. t and Ch. sp* | *Skeletonema sp. (CCAP1011/1B)* | (7) | *miTAG-v2_23481* | *99.8* | *188* |
| *Halomonas titanicae* | *T. p, T.w and Ch. sp.* | *Chaetoceros* sp. PLY617 | *This study* | *miTAG-v2_15166* | *98.9* | *2* |
| | | | | *miTAG-v2_6035* | *98.7* | *12* |
| | | | | *miTAG-v2_5294* | *98.6* | *2* |
| | | | | *miTAG-v2_13775* | *98.5* | *45* |
| | | | | *miTAG-v2_6036* | *98.4* | *1* |
| | | | | *miTAG-v2_6038* | *98.3* | *1* |
| | | | | *miTAG-v2_15164* | *97.7* | *1* |
| *Marinobacter adhaerens* | *T.w and Ch. sp.* | *Chaetoceros* sp. PLY617 | *This study* | *miTAG-v2_5840* | *100* | *8* |
| | | | | *miTAG-v2_22501* | *99* | *47* |

**Table S4. List of *Tara* stations the metabarcodes for the bacterial species described in Table S3 were identified.** Tara stations are coloured according to ocean regions (Mediterranean Sea, light blue; Red Sea, red; Indian Ocean, orange; South Pacific ocean, grey; South Atlantic, green, Arctic Ocean, purple; North Pacific, dark blue).

| *Thalassospira lohafexi* | *Maribacter dokdonensis* | *Marinobacter adhaerens* | *Halomonas titanicae* |
|---|---|---|---|
| *TARA_007* | *TARA_007* | *TARA_007* | *TARA_009* |
| *TARA_009* | *TARA_009* | *TARA_009* | *TARA_023* |
| *TARA_018* | *TARA_068* | *TARA_034* | *TARA_025* |
| *TARA_025* | *TARA_070* | *TARA_052* | *TARA_033* |
| *TARA_032* | *TARA_072* | *TARA_076* | *TARA_042* |
| *TARA_033* | *TARA_076* | *TARA_100* | *TARA_064* |
| *TARA_041* | *TARA_082* | *TARA_102* | *TARA_065* |
| *TARA_093* | *TARA_093* | *TARA_132* | *TARA_076* |
| *TARA_122* | *TARA_112* | | *TARA_094* |
| *TARA_168* | | | *TARA_122* |
| *TARA_173* | | | *TARA_128* |
| *TARA_188* | | | *TARA_132* |

**Table S5. List of metabarcodes identified in the TARA Ocean Barcode Atlas resembling 18S rDNA V9 region of diatom hosts effected by bacterial antagonists described in Table S3.** The query sequence and *Tara* Ocean barcode Identified for each hit, as well as details of % identity to the query sequence, and no. of reads detected are also given.

| Diatom species | NCBI identifier of query sequence | TARA Ocean Barcode Id. | Percentage identity (%) | No. of reads |
|---|---|---|---|---|
| *Thalassiosira pseudonana* | AY485452.1 | 18SV9-v2_1378; 18SV9v3_25785 | *100; 100* | *691; 660* |
| *Skeletonema marinoi* | AB948146.1 | 18SV9-v2_270 | 100 | *189089* |
| | | 18SV9-v2_157507 | 98.5 | 6 |
| | | 18SV9-v2_1614 | 98.5 | 90683 |
| | | 18SV9-v2_454417 | 98.5 | 3 |
| *Chaetoceros sp.* | KY852256.1 | 18SV9-v2_599 | *97.7* | *118967* |
| | | 18SV9-v2_468821 | *97.7* | *1* |
